# An integrative proteomics approach identifies tyrosine kinase KIT as a novel therapeutic target for SPINK1-positive prostate cancer

**DOI:** 10.1101/2023.07.24.550265

**Authors:** Nishat Manzar, Umar Khalid Khan, Ayush Goel, Shannon Carskadon, Nilesh Gupta, Nallasivam Palanisamy, Bushra Ateeq

## Abstract

Elevated Serine Peptidase Inhibitor, Kazal type 1 (SPINK1) levels in ∼10-25% of prostate cancer (PCa) patients associate with aggressive phenotype, for which there are limited treatment choices and dismal clinical outcomes. Using an integrative proteomics approach involving label-free phosphoproteome and proteome profiling, we delineated the downstream signaling pathways involved in SPINK1-mediated tumorigenesis in PCa, and identified tyrosine kinase KIT as a highly enriched kinase. Furthermore, high to moderate levels of KIT expression was detected in ∼85% of SPINK1-positive PCa specimens. KIT signaling regulates SPINK1-associated oncogenesis, and treatment with KIT inhibitor reduces tumor growth and distant metastases in preclinical mice models. Mechanistically, KIT signaling modulates WNT/β-catenin pathway and confers stemness-related features in PCa. Notably, inhibiting KIT signaling restores AR/REST levels, forming a feedback loop enabling SPINK1 repression. Overall, we uncover the role of KIT signaling downstream of SPINK1 in maintaining lineage plasticity and provide new treatment modalities for advanced-stage SPINK1-positive subtype.

## Introduction

Therapy resistance allied with disease recurrence is a pervasive problem in prostate cancer (PCa) patients. The standard of care for localized disease often involves surgical castration and/or androgen deprivation therapy (ADT) (1). Of these, majority of patients at some time point experience tumor relapse, leading to the emergence of castration-resistant prostate cancer (CRPC) (2, 3). Nearly one-fifth of metastatic CRPC patients succumb to small-cell neuroendocrine prostate cancer (NEPC) due to ADT-mediated lineage crisis (4, 5). Earlier, we have shown that AR-antagonists led to the upregulation of SPINK1 and genes associated with NEPC in CRPC mice models as well as in patients who underwent neoadjuvant-hormone therapy (6). The survival rate of these patients is less than a year, primarily due to its aggressiveness and limited treatment options (7).

High levels of Serine Peptidase Inhibitor, Kazal type 1 (SPINK1) represents the second-largest molecular subtype (∼10-25%) of PCa, associated with shorter biochemical recurrence and rapid progression to castration resistance (8-11). SPINK1 being a small secretory protein, acts in an autocrine/paracrine manner to promote tumor progression (12, 13). Furthermore, it has been linked with gastrointestinal-lineage signature genes aberrantly expressed in ∼30% of the CRPC cases owing to the activation of hepatocyte nuclear factors (HNF4G/HNF1A) transcriptional circuitry (14). In addition, genotoxic chemotherapy is known to trigger SPINK1 expression in the stromal cells, thereby promoting therapy resistance (15). Recently, we demonstrated that the Androgen Receptor (AR) and RE1-Silencing Transcription Factor (REST) function as transcriptional repressors of *SPINK1*, and AR antagonists release this repression resulting in its upregulation (6). Taken together, considering the functional significance of SPINK1 in PCa, there is a need to uncover its downstream unidentified signaling pathways, which may lead to the development of novel therapeutic strategies for this aggressive subtype.

Here, we delineate the downstream signaling pathways involved in SPINK1-mediated tumorigenicity by integrating the label-free phosphoproteome and proteome profiling data of SPINK1-positive PCa cells. Of all the enriched receptor tyrosine kinases, KIT receptor tyrosine kinase exhibits higher expression in NEPC compared to CRPC specimens. Additionally, we also examined KIT levels in PCa specimens, and ∼85% of SPINK1-positive PCa patients display high to moderate levels of KIT expression. We decipher that KIT signaling plays a crucial role in modulating the WNT/β-catenin pathway, and its inhibition restores the AR and REST levels, forming a feedback loop resulting in SPINK1 repression. Importantly, pharmacological inhibition of KIT resulted in the abrogation of oncogenic properties specifically associated with stemness phenotype, accompanied by reduced tumor growth and metastases. Collectively, we uncover the role of KIT signaling downstream of SPINK1, thus reinforcing its significance in modulating lineage plasticity and PCa progression.

## Results

### Global proteome and phosphoproteome profiling of SPINK1-positive prostate cancer

Dysregulated phosphorylation of proteins perturbs the activity of several biological pathways leading to tumorigenesis and metastases (16). Integrated proteomics studies offer an in-depth comprehension of cancer pathobiology and could untangle its complex circuitries. To decipher the signaling involved in SPINK1-mediated tumorigenesis, we performed label-free quantitative proteome and phosphoproteome profiling of small hairpin RNA-mediated *SPINK1* silenced (shSPINK1) and control small hairpin RNA Scrambled (shSCRM) 22RV1 cells (**Fig. S1**). Our integrated proteomics data revealed ∼4000 proteins and enriched ∼6000 phosphopeptides in each biological replicate. Of these, 367 proteins and 807 phosphopeptides were found to be significantly altered between shSCRM and shSPINK1 cells (**Fig. 1, A and B**). The differential enrichment analysis led to the identification of 190 downregulated proteins and 177 upregulated proteins (**Fig. 1C**).

**Figure 1.**
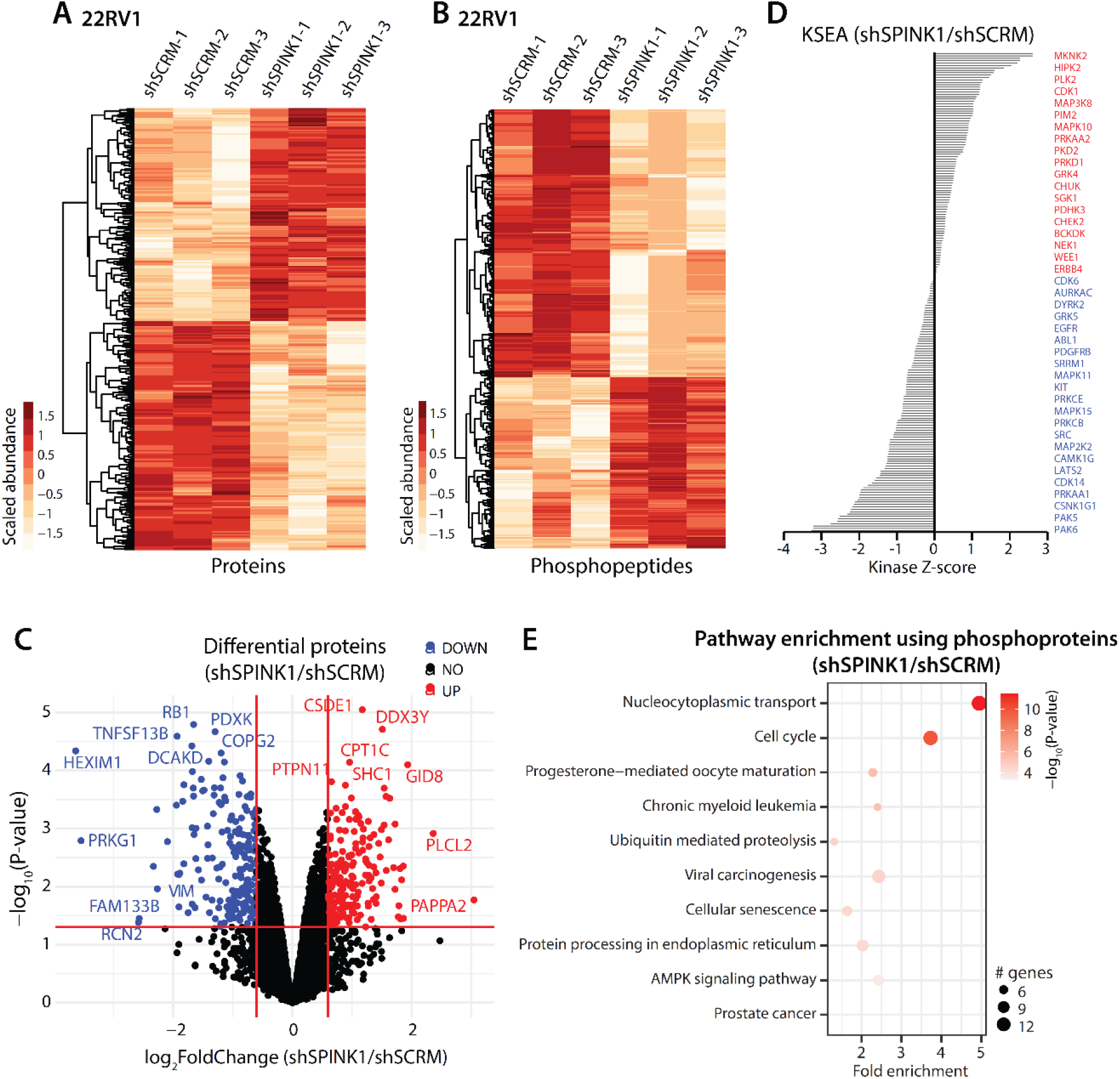
Global quantitative proteome and phosphoproteome profiling of the SPINK1-knockdown prostate cancer cells. (**A**) Heatmap depicting the Z-scaled abundance of significant proteins in 22RV1-shSCRM and 22RV1-shSPINK1 cells. (**B**) Same as in (A) except for significantly enriched phosphoproteins in 22RV1-shSCRM and 22RV1-shSPINK1 cells. Each sample was analyzed in biological triplicates. **(C**) Volcano plot showing the differentially enriched proteins in 22RV1-shSPINK1 versus 22RV1-shSCRM cells; blue dots are downregulated, and red dots are upregulated proteins in shSPINK1, whereas black dots signify no change. (**D**) Kinase-substrate enrichment analysis of the phosphopeptides enriched in 22RV1-shSPINK1 versus 22RV1-shSCRM cells. Identified kinases are plotted with their respective z-scores; blue kinases have negative z-score, and red kinases have positive z-score in 22RV1-shSPINK1 versus 22RV1-shSCRM cells. A negative z-score corresponds to a collective dephosphorylation of the kinase’s substrates; the inverse is true for a positive score. (**E**) Pathway enrichment analysis for differential phosphoproteins in 22RV1-shSPINK1 versus 22RV1-shSCRM cells using pathfindR; size of the dot represents number of genes, and the color symbolizes the -log_10_(P-value) of the enriched pathway.

Also, 492 downregulated phosphopeptides and 315 upregulated phosphopeptides were noted in 22RV1-shSPINK1 versus 22RV1-shSCRM cells (**Fig. S2A**). Since deregulated kinases have been associated with cancer and kinase-targeting drug such as imatinib has been a great success in multiple cancer types (17, 18), we examined the kinases involved in this differential phosphorylation of proteins using Kinase-Substrate Enrichment Analysis (KSEA) (19). We identified ∼200 kinases to be enriched in the kinase-substrate relationship, and decreased activity of 108 kinases, including MET, CSF1R, FLT3, KIT, INSR, PDGFRB, ERBB2, and EGFR, was noticed in SPINK1 ablated cells relative to the control (**Fig. 1D**). Furthermore, to identify the biological pathways governed by the dysregulated proteins and phosphopeptides, we performed pathway enrichment analysis using pathfindR and DAVID functional annotation tools (20, 21). Notably, cell cycle, ubiquitin-mediated proteolysis, protein processing in endoplasmic reticulum, AMPK signaling, and prostate cancer appeared as few of the top ten significantly altered pathways with SPINK1 silencing (**Fig. 1E and S2, B and C**). These findings reveal that SPINK1 engages distinct kinases to direct the activation or deactivation of downstream signaling pathways in promoting tumorigenesis and stemness in prostate cancer.

### Sustained receptor tyrosine kinase KIT signaling and SPINK1 drive neuroendocrine prostate cancer

ADT-induced SPINK1 plays a crucial role in governing stemness as well as in the maintenance of neuroendocrine phenotype (6). To explore the SPINK1-activated receptor tyrosine kinase in advanced-stage PCa, we examined the expression of different kinases in publicly available NEPC datasets (**Fig. 2A and S3, A-G**). Interestingly, higher expression of *KIT* and *INSR* kinases in NEPC (N=15) compared to CRPC specimens (N=34) was observed in the Beltran cohort (**Fig. 2A and S3G**). Similarly, high expression of *KIT* was noted in small-cell NEPC (SCNC) (N=12) compared to metastatic CRPC (N=107) specimens in the Aggarwal *et al.* dataset (4) (**Fig. 2B**). However, no significant change in *INSR* expression between metastatic CRPC and small-cell NEPC specimens was detected in Aggarwal *et al.* dataset (**fig. S3H**). Next, to examine a plausible link between KIT and androgen signaling, we analysed the association of *KIT* and AR signaling scores in the metastatic prostate adenocarcinoma (SU2C/PCF Dream Team) cohort (N=264). Similar to *SPINK1* expression, *KIT* also displayed an inverse association with AR signaling (**Fig. 2C and S4A**). Additionally, a positive correlation between *KIT* and NEPC gene signature was seen in metastatic CRPC, concurring with a positive association of SPINK1 and NEPC score (**Fig. 2D and S4B).** Notably, no significant association of INSR with either AR signaling or NEPC score was detected in CRPC specimens (**fig. S4, C and D**). Next, we performed immunohistochemical (IHC) staining for the expression of KIT and SPINK1 in the tissue microarray (TMA), and found that ∼85% of SPINK1-positive PCa specimens (N=49) exhibited high to medium levels of KIT expression (**Fig. 2, E and F and S4E**). Subsequently, we examined the *SPINK1* and *KIT* expression in NEPC patients derived organoids (GSE112786), and found that SPINK1-positive NEPC organoids displayed higher levels of *KIT* expression corroborating with our cell lines data (**Fig. S4F**). We also checked *SPINK1* and *KIT* expression in advanced PCa specimens who underwent ADT for ∼22 weeks (GSE48403), and noticed high levels of *SPINK1* and *KIT* in post-ADT patients’ specimens relative to pre-ADT (**Fig. 2G and S4G**). Collectively, these results highlight the association of receptor tyrosine kinase KIT signaling and SPINK1 in androgen-independent prostatic tumors.

**Figure 2.**
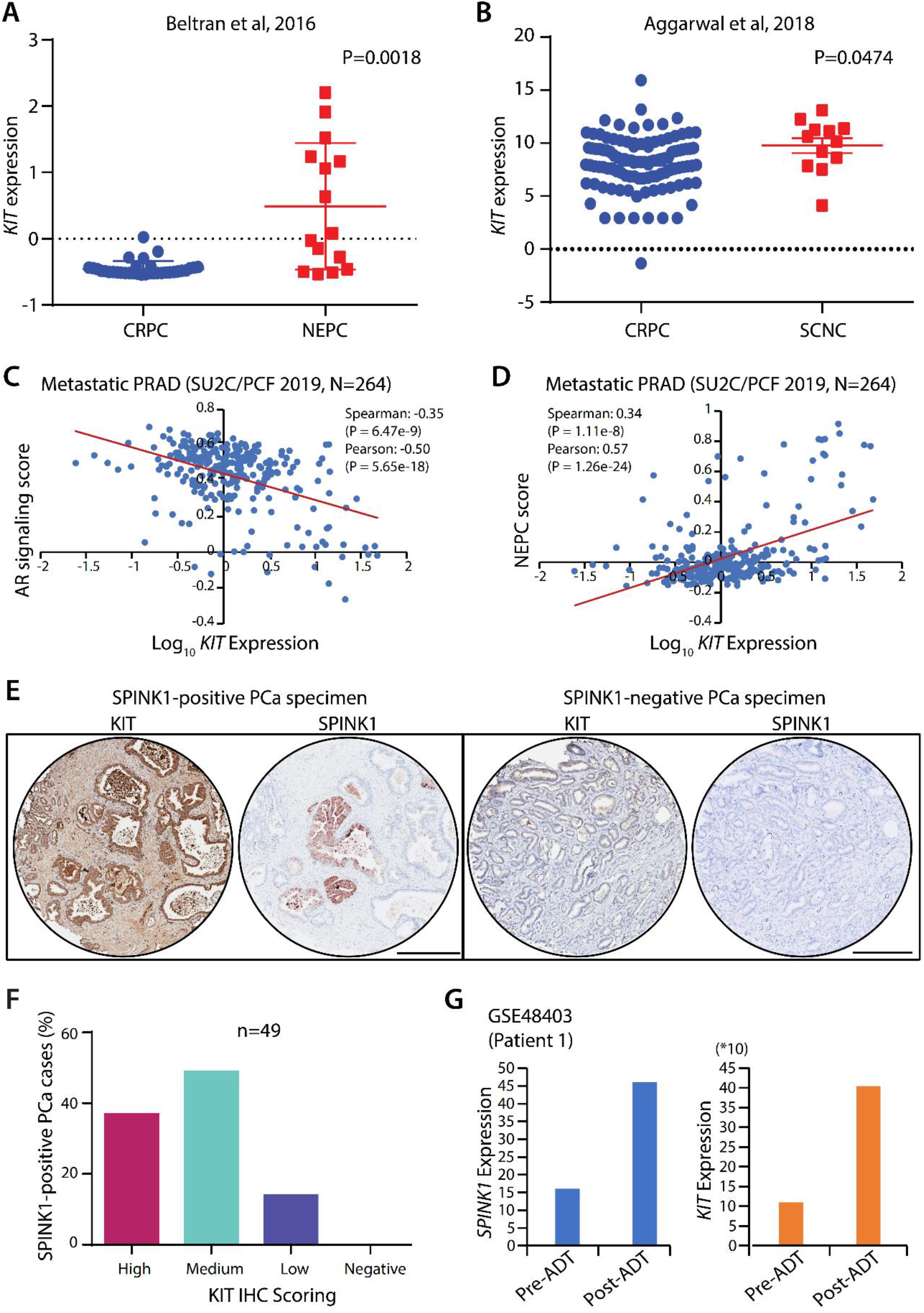
KIT tyrosine kinase is highly expressed in androgen-independent prostatic tumors. (**A**) *KIT* mRNA expression in Beltran et al., 2016 dataset. P-value was calculated using Unpaired Student’s t-test with Welch’s correction. (**B**) Same as (A) except for Aggarwal et al., 2018 dataset. (**C**) Scatter plot showing correlation of *KIT* mRNA expression (FPKM polyA) (log_10_(value+1)) with AR signaling score in metastatic prostate adenocarcinoma (SU2C/PCF 2019) dataset. Coefficient values for both Spearman and Pearson’s correlation along with the P-value are depicted in the plot. (**D**) Same as in (C) except for the correlation of *KIT* mRNA expression with NEPC score. (**E**) Micrographs representing immunohistochemical (IHC) staining for KIT and SPINK1 expression in tissue microarrays of prostate cancer specimens, SPINK1-positive patient (left) and SPINK1-negative patient (right). Scale bar represents 300µm. (**F**) Bar plot depicting the percentage of SPINK1-positive patients’ specimens for KIT intensity (high/medium/low/negative). (**G**) Bar plots showing *SPINK1* and *KIT* expression in 22 weeks ADT treated PCa patient with Gleason score-8 and TNM stage-3b.

### Pharmacological inhibition of KIT signaling perturbs SPINK1-mediated tumorigenesis

Having established the connection between receptor tyrosine kinase KIT signaling and SPINK1, we next determine the protein level expression of KIT and SPINK1 across different PCa and benign prostate epithelial cells. To our interest, both the proteins exhibited high expression in NCIH660, an epithelial neuroendocrine cell line, followed by a moderate expression in 22RV1, a castrate-resistant cell line (**Fig. S5, A and B**). Next, to investigate the role of KIT signaling in SPINK1-mediated tumorigenesis, we treated 22RV1 cells with varying concentrations of KIT tyrosine kinase inhibitor (KITi), and a concentration-dependent inhibition of cell proliferation and viability of KITi treated 22RV1 cells was observed (**Fig. 3, A and B**). Notably, half-maximal inhibitory concentration (IC50) of KITi for 22RV1 cells was 16.9µM with ∼50% reduction in cell viability (**Fig. S5C**). To examine the effect of KIT kinase in anchorage-independent growth, foci formation ability of 22RV1 cells was performed in presence and absence of KITi, and an 80% reduction in the foci formation ability of 22RV1 cells was noticed with a sub-IC50 concentration (10µM) of KITi (**Fig. 3C**).

**Figure 3.**
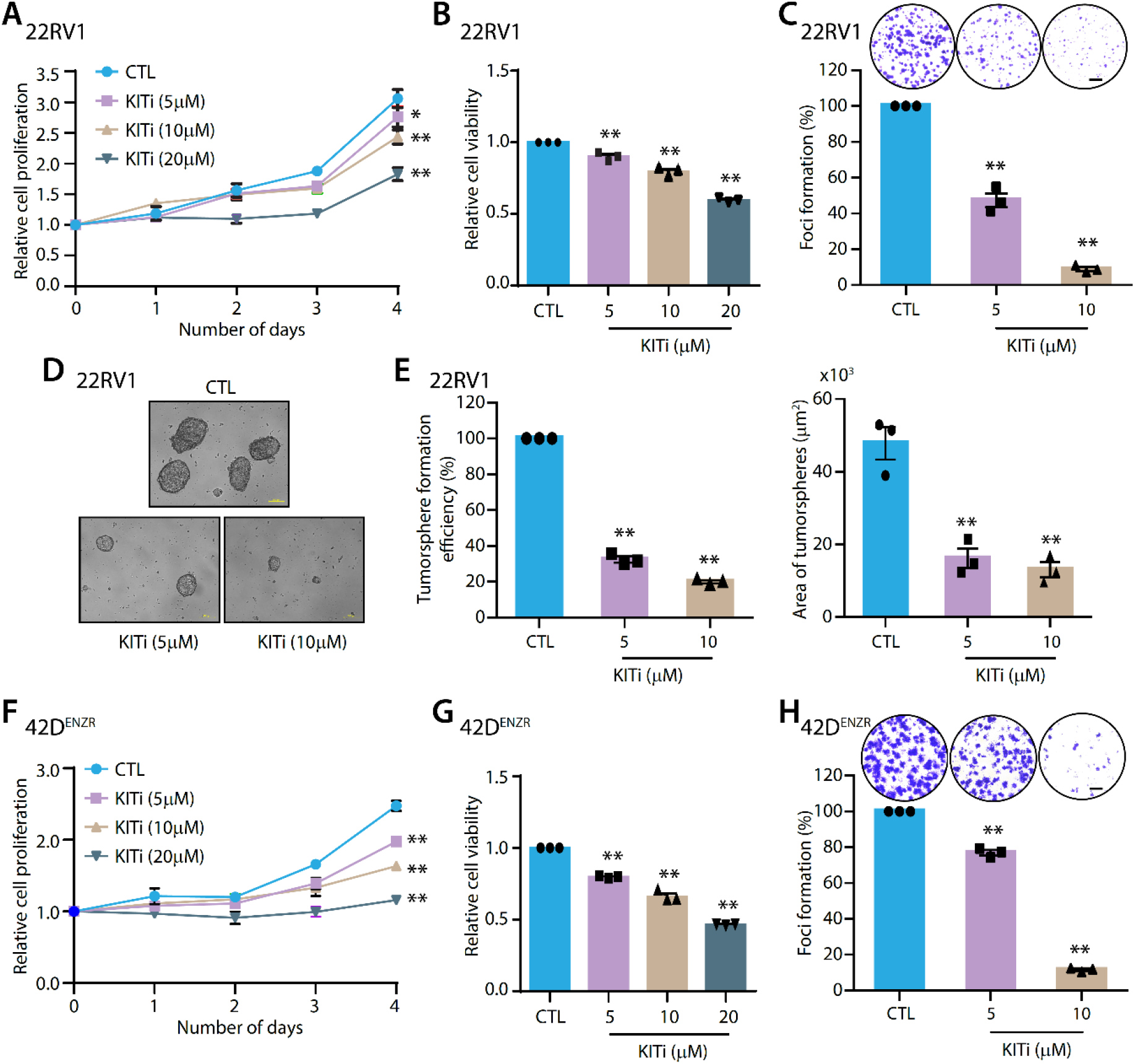
KIT tyrosine kinase inhibition attenuates oncogenic characteristics. (**A**) Line plot representing relative cell proliferation of 22RV1 cells along with different concentrations of KIT tyrosine kinase inhibitor, KITi and control (CTL). (**B**) Same as in (A), except for bar plot showing relative cell viability. (**C**) Representative images for foci of 22RV1 cells in the respective conditions (Top). Bar plot showing percent foci formation efficiency of 22RV1 cells treated with indicated concentrations of KITi and CTL. Scale bar represents 1000µm. (**D**) Representative images for 22RV1 tumorspheres treated with KITi/CTL. Scale bar denotes 100µm. (**E**) Bar plots showing percent tumorsphere formation efficiency (left) and mean area of 22RV1 tumorspheres (right) treated with indicated concentrations of KITi and CTL. (**F**) Same as in (A) except for 42D^ENZR^ cells. (**G**) Same as in (B) except for 42D^ENZR^ cells. (**H**) Representative images for foci of 42D^ENZR^ cells in the respective conditions (Top). Bar plot showing percent foci formation of 42D^ENZR^ cells treated with the indicated concentrations of KITi and CTL. Scale bar represents 1000µm. Each experiment was performed in biological triplicates (N=3); the bar represents mean ± SEM, and each dot represents individual value. Statistical significance was calculated using Two-way or One-way ANOVA followed by Dunnett’s multiple comparison test. P-value: *<0.05 and **<0.001.

To evaluate the significance of KIT signaling in conferring stemness and self-renewal capacity, tumorsphere formation assay was performed and ∼70-80% reduction in tumorsphere formation efficiency and mean area of 22RV1 tumorspheres treated with KITi was noted compared to control (**Fig. 3, D and E**). Further, to investigate the effect of KIT signaling inhibition in therapy-resistant PCa, the expression of *SPINK1* and *KIT* was first examined in 16D^CRPC^ treated with Enzalutamide for 10 days and enzalutamide-resistant 42D^ENZR^ cells (22). Notably, an increase in *SPINK1* expression in castrate-resistant 16D^CRPC^ was observed, while in enzalutamide-resistant 42D^ENZR^ cells both *SPINK1* and *KIT* were elevated (**Fig. S5D**). Moreover, marked reduction in cell proliferation and viability of 42D^ENZR^ cells with increasing concentrations of KITi was noted (**Fig. 3, F and G**). Alongside, KITi (10µM) treatment also led to a robust decrease in the foci formation ability of 42D^ENZR^ cells (**Fig. 3H**) supporting the probable function of KIT signaling in imparting therapy resistance in conjunction with high SPINK1 levels. These results depict the central role of KIT signaling in governing oncogenic attributes of SPINK1-positive PCa.

### Pharmacological inhibition of KIT signaling mitigates SPINK1-mediated cancer stemness

KIT has been characterized as one of the important markers of prostate stem cells, and is known to play a key role in the organogenesis of prostate gland (23). In PCa, KIT is known to drive disease aggressivity through the cancer stem-like cells (CSCs) phenotype and resistance to tyrosine kinase inhibitor (24). To evaluate the role of KIT signaling in maintaining cancer-associated stemness, RNA was isolated from the tumor spheres generated in Figure 3D. The expression of markers associated with cellular reprogramming was examined, and a notable decrease in the expression of *MYC*, *OCT4*, *SOX2*, *TET1*, and *AURKA* was noticed in KITi- treated tumorspheres compared to control (**Fig. 4A**).

**Figure 4.**
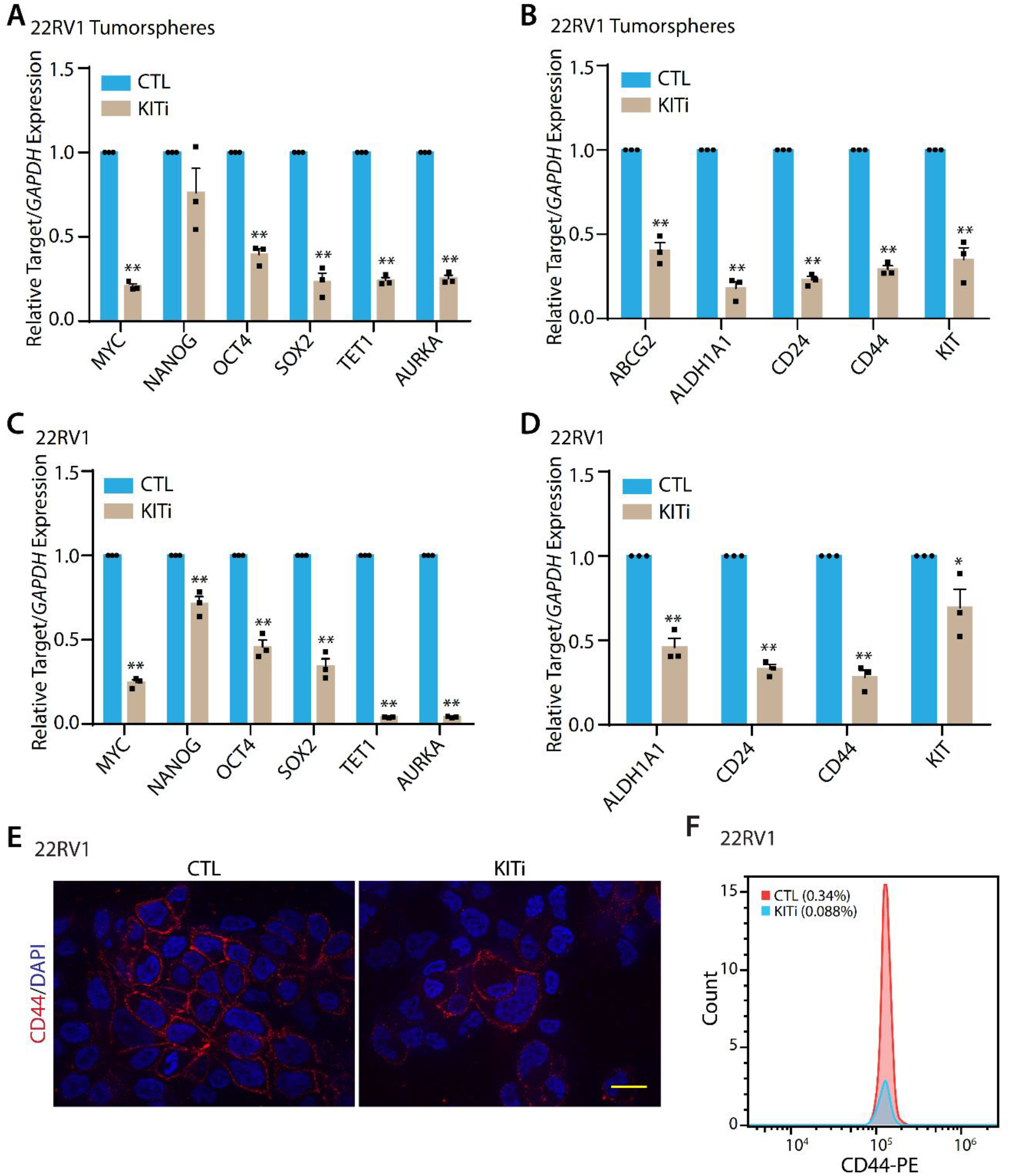
KIT tyrosine kinase inhibition mitigates stemness-related features. (**A**) Bar plot showing quantitative-PCR (qPCR) data for the relative expression of stemness-associated genes in KITi (10µM)/CTL-treated 22RV1 tumorspheres. (**B**) Same as in (A), except for relative expression of stem cell surface markers (**C**) Same as in (A), except for KITi/CTL-treated 22RV1 cells. (**D**) Same as (B) except for KITi/CTL-treated 22RV1 cells. (**E**) Micrographs representing immunostaining for CD44 in KITi/CTL-treated 22RV1 cells. Scale bar represents 10µm. (**F**) Histogram depicting flow cytometry analysis of CD44 staining in KITi/CTL-treated 22RV1 cells. Each experiment was performed in biological triplicates (N=3); the bar represents mean ± SEM and each dot represents individual value. Statistical significance was calculated using Two-way ANOVA followed by Sidak’s multiple comparison test. P-value: *<0.05 and **<0.001.

Furthermore, KITi-treated tumorspheres showed a robust decrease in the expression of stem cell surface markers, namely, *CD24*, *CD44,* and *KIT*/*CD117* (**Fig. 4B**). In agreement with tumorsphere data, a similar trend of reduced expression of stemness associated genes and cell surface markers in KITi treated 22RV1 cells was noticed (**Fig. 4, C and D**). To consolidate the changes at the gene expression level of cell surface stem cell markers, we also examined the localization and surface expression of CD44, and ∼75% reduction in the cell surface levels of CD44 was recorded upon treatment with KITi compared to control (**Fig. 4, E and F**). In addition, KIT signaling inhibition in androgen-deprived LNCaP cells (LNCaP-AI), which express higher SPINK1, also demonstrated a modest decrease in the expression of stem cell surface markers (**Fig. S5E**). These findings emphasize the role of KIT signaling in maintaining stemness in the context of SPINK1-positive cancers.

### Inhibition of KIT diminished SPINK1-mediated osteolytic bone metastases

The role of KIT signaling has been well-defined in the progression of gastrointestinal stromal tumors (25); however, its allies in prostate tumorigenesis remain elusive. To address this question, we generated Doxycycline (Dox) inducible KIT overexpressing 22RV1 cells, namely, 22RV1-KIT (**Fig. S6A**), and KIT signaling inhibition via a decrease in phosphorylation levels of KIT using KITi was confirmed (**Fig. 5A**). To identify the KIT interacting partners, we performed immunoprecipitation coupled with tandem mass spectrometry (IP-MS) and identified 306 proteins exclusive to KIT pull-down assay (**Fig. 5B**). Poly [ADP-ribose] polymerase 1 (PARP-1), histones, heterogeneous nuclear ribonucleoproteins (hnRNPs), the 40S and 60S ribosomal proteins (40S/60S RPs), proteasome subunit α and β (PSMAs/PSMBs) were some of the proteins enriched in KIT IP-MS (**Fig. 5B and Table S1)**. Biological processes enrichment using KIT interacting partners (N=306) discovered multiple significant pathways (**Fig. 5C**). Surprisingly, the majority of these processes regulated by KIT interactome show similarity to the processes associated with SPINK1 downregulated proteins, for instance, translation, mRNA processing and RNA splicing (**Fig. S6B**). Furthermore, 33 proteins were shared among KIT-interacting proteins and SPINK1-downregulated proteins, while 19 proteins were seen as common between KIT interactome and SPINK1 upregulated (**Fig. S6C**). These findings imply the involvement of KIT signaling in the regulation of biological processes, such as mRNA processing, splicing and translation in SPINK1-positive PCa.

**Figure 5.**
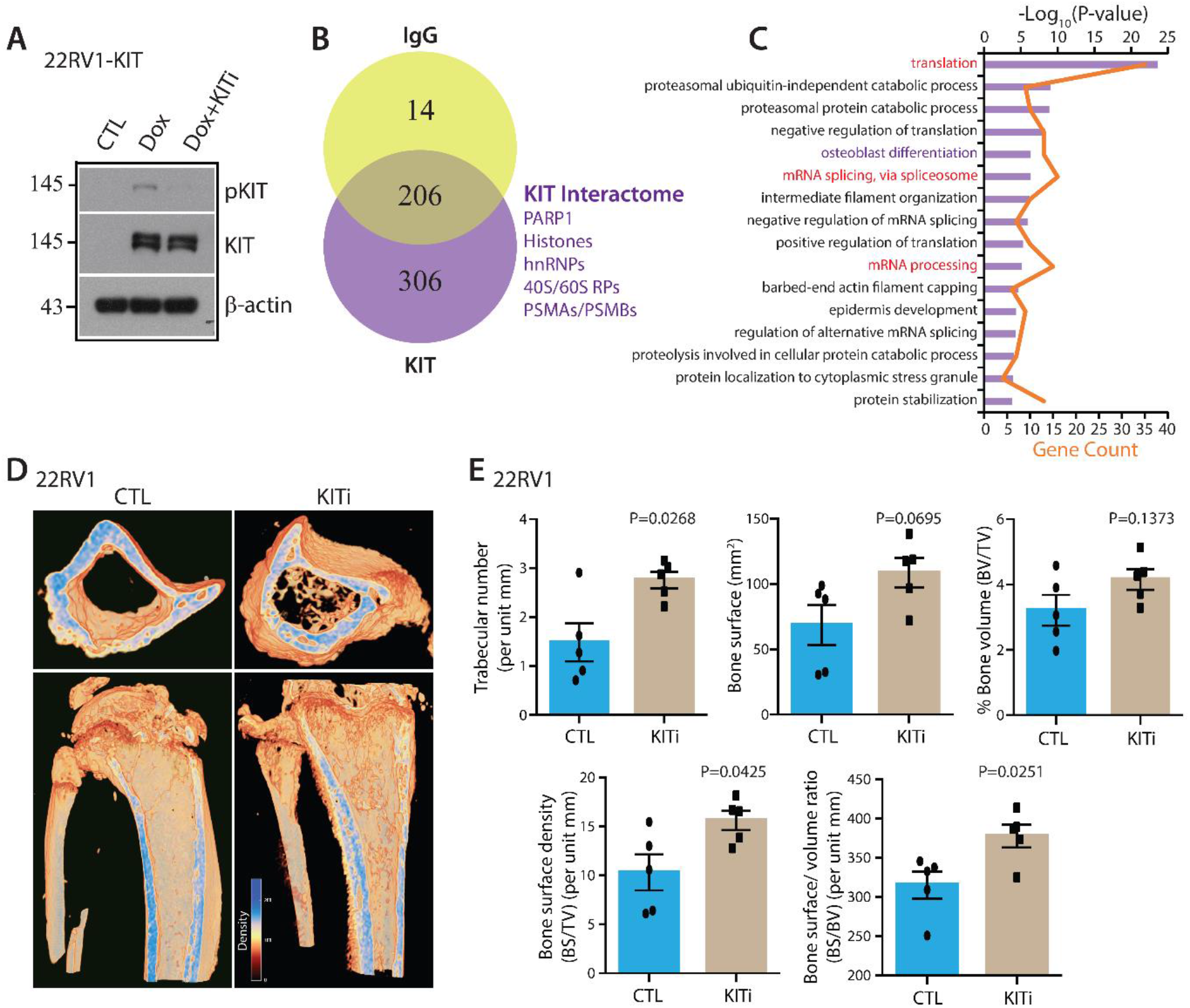
KIT tyrosine kinase inhibition limits metastatic bone lesions. (**A**) Immunoblot showing phosphorylated-KIT (pKIT) and KIT levels in 22RV1-KIT cells induced with Doxycycline (Dox) and treated with KITi. β-actin was used as the loading control. (**B**) Venn diagram depicting common and specific proteins between IgG and KIT pulldown in Dox-induced 22RV1-KIT cells. A few KIT interacting partners are also labelled. (**C**) DAVID functional annotation analysis showing enriched biological processes using KIT interactome (N=306). The bar denotes -log_10_(P-value), and the line represents gene count. (**D**) Micrographs representing micro-CT analysis of proximal horizontal (top) and longitudinal cross-sectional view (bottom) of the tibiae from mice with 22RV1 intratibial implantation and administered with KITi (50mg/kg)/CTL via oral gavage; (each group, N=5). (**E**) Bar plots showing bone morphometric parameters in the tibiae, same as shown in (D). Bar represents mean ± SEM and each dot represents the individual value. Statistical significance was calculated using Unpaired Student’s t-test with Welch’s correction. P-value: *<0.05 and **<0.001.

Interestingly, KIT interactome pathway analysis highlighted osteoblast differentiation as one of the significant biological processes (Fig. 5C). In analogy, KIT is constitutively expressed on the human osteoclasts, and plays key role in bone resorption and remodeling (26). Since 22RV1 tumors associated bone metastases demonstrate both osteolytic and osteoblastic features (27), we aimed to examine the effect of KIT signaling using an experimental bone metastasis model. Towards this, 22RV1 cells were implanted via intramedullary injection in the tibia of NOD/SCID mice, and administered them with KITi (pexidartinib, 50mg/kg) or vehicle control (CTL) orally for three weeks. Bone metastases in the tibiae was monitored by X-ray, and the characteristics and extent of bone lesions were examined by micro-computed tomography (µCT) (**Fig. S7, A and B and 5D**). Notably, a significant increase in the number of trabeculae suggestive of more cancellous bone remodelling was observed in the mice treated with KITi (**Fig. 5, D and E**). Furthermore, assessment of the proximal epiphysis and metaphysis regions revealed moderate changes in the bone morphometric parameters, such as bone surface density and bone surface/volume ratio with KIT inhibition (**Fig. 5E**). These findings employ the role of KIT signaling in osteolytic PCa bone metastases.

### Targeting KIT signaling restores AR/REST axis by impairing WNT signaling

The CD44 stem cell marker has long been defined as a target of WNT/β-catenin pathway in the intestinal tumorigenesis (28). In our IP-MS data, KIT showed direct interaction with β- catenin with 9 peptide spectral matches (Table S1). Also, KIT signaling inhibition resulted in diminished stemness and reduced CD44 levels **(Fig. 4, E and F)**; therefore, we conjectured its effect on WNT/β-catenin pathway. Notably, a rampant decrease in the expression of genes associated with WNT signaling, such as *CTNNB1*, *TCF7L2*, *SOX9,* and *AXIN2,* was found in KITi treated 22RV1 tumorspheres as well as cells (**Fig. 6, A and B**). Similar results were obtained in androgen-deprived LNCaP-AI cells treated with KITi (**Fig. S8A**). Likewise, KITi treatment ensued a concentration-dependent decrease in the protein levels of β-catenin, CD44 and KIT relative to the control (**Fig. 6C**). In contrast, inducible KIT overexpression in 22RV1 cells demonstrated high levels of β-catenin compared to the control (**Fig. S8B**), proving the notion that KIT signaling regulates stemness associated β-catenin pathway in prostate tumorigenesis. Furthermore, KITi treatment led to reduced phosphorylation of ERK and AKT (**Fig. S8C**), which is a prototypic readout of KIT tyrosine kinase inhibition. On another note, the PI3K-Akt signaling pathway is known to harness ubiquitin-mediated proteasomal pathway to degrade AR (29). Interestingly, ’ubiquitin mediated proteolysis’ and ’proteasomal protein catabolic process’ showed up as the top significant pathways enriched in the SPINK1 regulated phosphoproteins and KIT interacting proteins (**Fig. 1E and 5C**).

**Figure 6.**
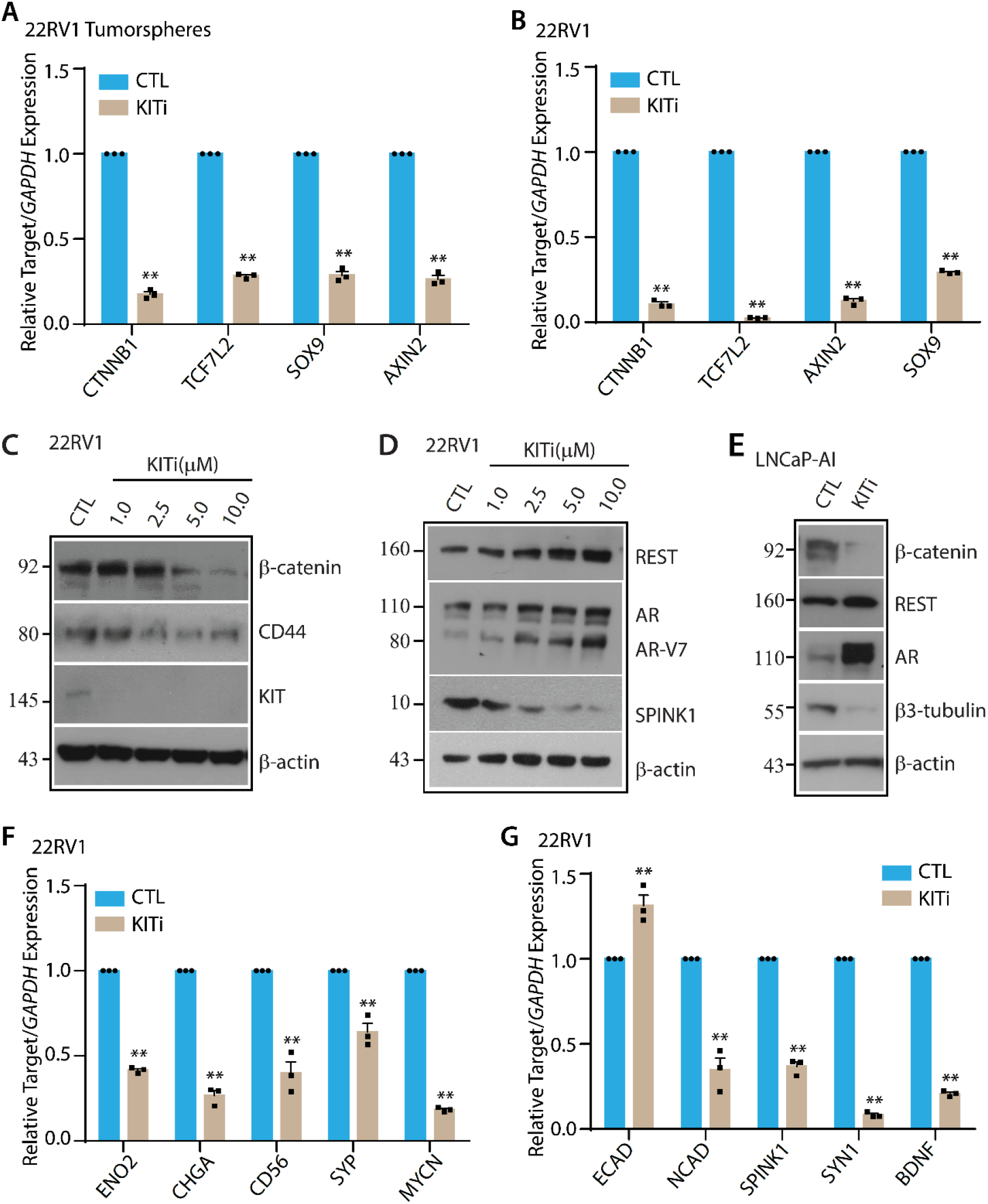
KIT tyrosine kinase inhibition dysregulates the WNT/β-catenin pathway and reinstates AR/REST axis. (**A**) Bar plot depicting qPCR data for the relative expression of WNT/β-catenin pathway genes in KITi (10µM)/CTL-treated 22RV1 tumorspheres. (**B**) Same as in (A) except for KITi/CTL-treated 22RV1 cells. (**C**) Immunoblot showing the expression level of β-catenin, CD44 and KIT in 22RV1 cells treated with indicated concentrations of KITi and CTL. β-actin was used as the loading control. (**D**) Immunoblot showing the expression level of REST, AR/AR-V7 and SPINK1 using same cells as in (D). (**E**) Immunoblot showing expression level of β-catenin, REST, AR and β3-tubulin in KITi (10µM)/CTL-treated LNCaP-AI cells. β-actin was used as the loading control. (**F**) Bar plot representing qPCR data for the relative expression of NEPC markers using the same cells as in (B). (**G**) Bar plot showing qPCR data for the relative expression of EMT markers, *SPINK1* and REST target genes using the same cells as in (B). Each experiment was performed in biological triplicates (N=3); bar represents mean ± SEM and each dot represents individual value. Statistical significance was calculated using Two-way ANOVA followed by Sidak’s multiple comparison test. P-value: *<0.05 and **<0.001.

Taking cues from these results, we interrogated the effect of KIT signaling inhibition on AR levels, and a concentration-dependent rise in AR and its corepressor REST protein levels was observed upon KITi treatment (**Fig. 6D**). In line with our previous study (6), increase in AR and REST upon KITi treatment led to a concomitant decrease in SPINK1 levels (**Fig. 6D**), forming a feedback loop. Similarly, KIT signaling inhibition led to AR/REST restoration in LNCaP-AI cells, which in turn hampered neuroendocrine phenotype as marked by a decrease in the β3-tubulin levels (**Fig. 6E**). Moreover, a remarkable reduction in NEPC markers (*ENO2*, *CHGA*, *CD56,* and *NMYC*), REST target genes (*SYN1* and *BDNF*) was noted in KITi treated 22RV1 cells and tumorspheres compared to control (**Fig. 6, F and G and S8D**). Also, KIT signaling inhibition showed a decrease in *NCAM*, a mesenchymal marker, along with a modest increase in epithelial markers (*ECAD* and *EPCAM*) (**Fig. 6G and S8D**), suggesting its role in maintaining cellular plasticity. Collectively, these results lay down a novel mechanism wherein KIT signaling regulates the stability of AR/REST, which in turn governs the transition to neuroendocrine phenotype.

### Pharmacological inhibition of KIT signaling attenuates tumor growth and metastases

To understand the therapeutic utility of KIT inhibitor in SPINK1-mediated PCa progression, 22RV1 cells were subcutaneously implanted in the flank region of NOD/SCID mice and tumor burden was monitored. The mice were randomized into two groups once the mean tumor volume reached ∼75mm^3^ and mice were administered with KITi (50mg/kg) or vehicle control (CTL) orally for three weeks. Mice treated with KITi exhibited diminished tumor growth and ∼50% reduction in tumor volume. (**Fig. 7, A and B**). Importantly, KITi treatment showed no adverse effect, as evident by no change in mice body weight between the two groups (**Fig. 7C**). Next, we sought to determine the impact of KITi on tumor cell stemness and proliferation by performing immunohistochemistry (IHC) staining for CD44 and Ki67 using tumor xenografts. KITi treatment significantly decreased levels of CD44 and Ki67 relative to the CTL group (**Fig. 7, D and E**). Subsequently, we also examined the effect of KITi on spontaneous distant metastases and checked the expression of human-specific *Alu* repeats using genomic DNA isolated from the bone marrow and lungs of the mice of both treatment and control groups (30). KITi-treated mice displayed a reduced number of cells metastasized to distant organs, such as bone and lungs compared the CTL group (**Fig. 7, F and G**). These findings summarize the efficacy of pharmacological inhibition of KIT signaling in abrogating SPINK1-mediated prostate tumorigenesis.

**Figure 7.**
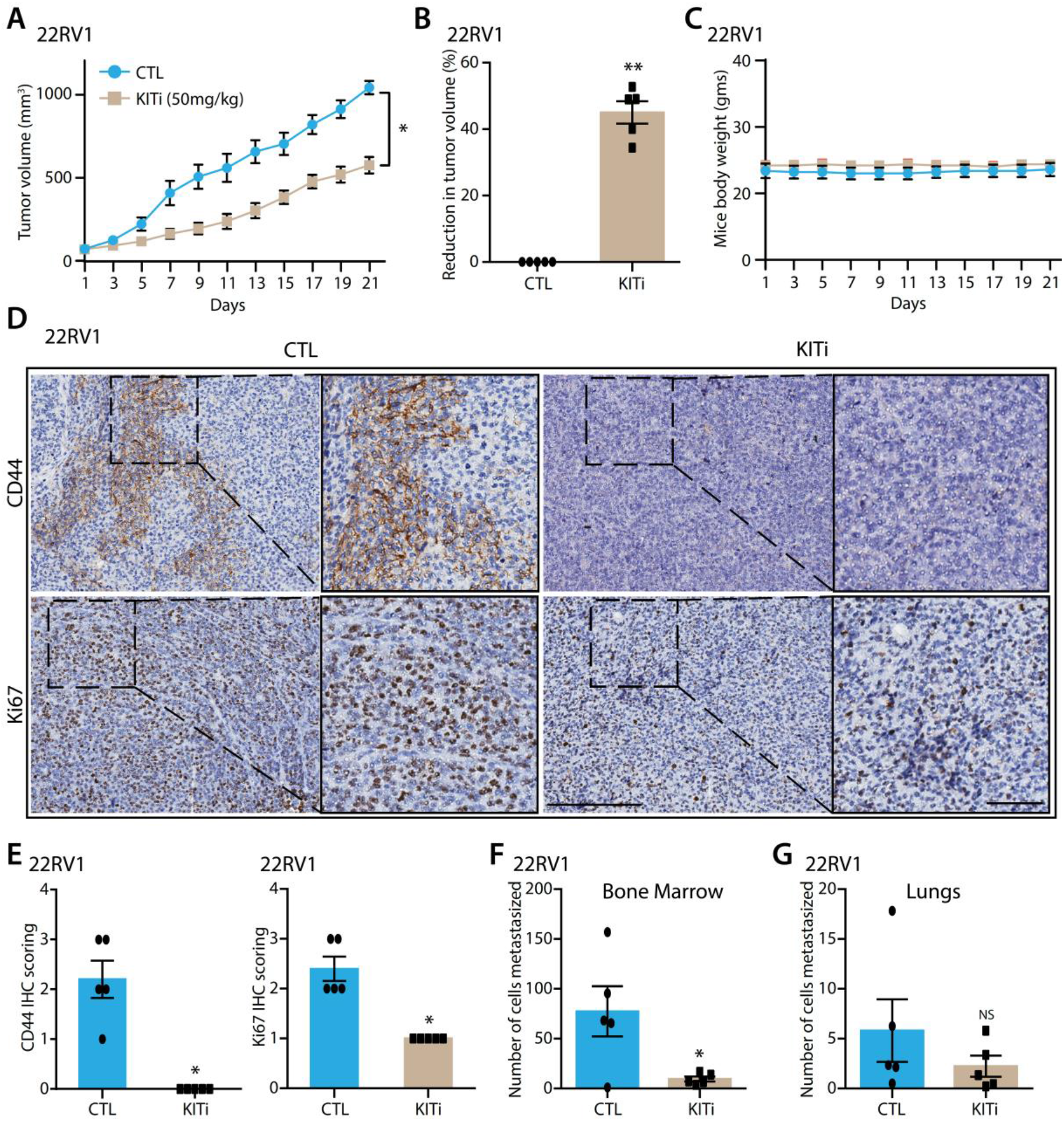
KIT tyrosine kinase inhibition abrogates SPINK1-positive tumor growth and metastases. (**A**) Line plot representing tumor volume of 22RV1 cells subcutaneously implanted in NOD/SCID mice and administered with KITi (50mg/kg)/CTL via oral gavage (each group, N=5). (**B**) Bar plot showing the relative percent reduction in tumor volume. (**C**) Same as in (A) except for the plot depicting mice’s body weight. (**D**) Micrographs representing IHC staining for CD44 and Ki67 in tumor sections from xenografted mice as shown in (A) Scale bar represents 200µm and 50µm (for insets). (**E**) Bar plots showing quantification of IHC staining for CD44 and Ki67. Twenty random fields were quantified for each group. (**F**) Bar plots representing the number of cells metastasized to the bone marrow in xenografted mice, as shown in (A). (**G**) Same as in (F) except for the number of cells metastasized to the lungs. Bar represents mean ± SEM and each dot represents individual value. Statistical significance was calculated using Unpaired Student’s t-test. P- value: *<0.05, **<0.001 and NS = non-significant.

## Discussion

SPINK1 molecular subtype has been associated with more aggressive disease and overall poor clinical outcomes (8-11). Earlier, we have shown that AR-antagonists alleviate AR- and its corepressor, REST-mediated transcriptional repression of *SPINK1* in PCa. Moreover, lineage reprogramming factor SOX2 transactivates *SPINK1* during neuroendocrine transdifferentiation, leading to its upregulation (6, 31). A recent study showed that SPINK1 mitigates radiation-induced DNA damage by upregulating EGFR- and Nrf2-dependent antioxidant responses, subsequently leading to cancer radioresistance (32). Previously, Ateeq and colleagues demonstrated that SPINK1 interacts with epidermal growth factor receptor (EGFR) owing to its structural homology with EGF, and activates downstream signaling cascade, nonetheless they conjectured that EGFR-independent pathways may also be involved in SPINK1-mediated oncogenic effects (13).

This study deciphers the activation of EGFR-dependent as well as -independent signaling pathways in SPINK1-positive PCa. Since the EGFR-targeted therapies have demonstrated limited success in metastatic CRPC patients (33, 34), our prime focus was to interrogate the receptor tyrosine kinases, which might function distinctly in androgen- independent prostatic tumors. Different members of the type III receptor tyrosine kinase family, including CSF1R, KIT, FLT3, and PDGFR, exhibited reduced kinase activity upon silencing of SPINK1. Consistent with the impact of ADT on SPINK1, KIT also displayed higher expression post-ADT and an inverse relation with AR signaling, implicating an association of KIT and SPINK1 in CRPC progression.

Since therapeutic targeting of KIT signaling leads to ∼50% reduction in cell proliferation, a pronounced decrease in foci formation, and tumorsphere formation ability, we reasoned the predominant role of KIT in imparting stemness-related attributes, which was validated via the change in the stemness-associated transcription factors and cell surface markers. In line with our findings, KIT is recently shown as a PCa stem cell marker (24). Besides, the activation and therapeutic potential of KIT ligand-induced canonical KIT signaling has recently been established in NEPC (35). These reports confirm the critical role of KIT in lineage plasticity, however the exact mechanism of KIT signaling and its downstream players remain unexplored in PCa progression.

Our integrated proteome data from SPINK1 knockdown and KIT interactome stipulated mRNA processing, splicing, and translation as some of the commonly altered biological processes. A closer view of the KIT interactome revealed PARP-1, DNA-dependent protein kinase, catalytic subunit (PRKDC), many histones, and hnRNPs as its direct binding partner (Table S1). Concurrently, histones (H1, H2A, H2B, and H4), hnRNPs (HNRNPC, HNRNPK, HNRNPM, HNRNPU), and PRKDC were also reported as PARP1 interacting partners by affinity purification mass spectrometry (36). Also, several studies have proven the canonical and non-canonical function of PARP1 in RNA processing, splicing, and translation (37, 38). These findings suggest a plausible role of KIT signaling in tuning RNA metabolism in SPINK1-positive PCa.

We have unraveled β-catenin as a KIT interacting protein, and targeting KIT signaling disrupted the WNT/β-catenin pathway. A previous study in mast cell leukemia has reported similar results, where activated KIT directly interacts with β-catenin and leads to its tyrosine phosphorylation, which triggers its nuclear localization and enhanced transcriptional activity (39). Moreover, elevated levels of β-catenin nuclear localization has been observed in CRPC bone metastases (40). Our experimental bone metastases mice data also highlighted the role of KIT signaling in bone resorption, where its inhibition resulted in cancellous bone remodeling. These findings demonstrated that the SPINK1-activated KIT kinase modulates β-catenin stability and transactivates target genes in CRPC. Inhibition of KIT signaling concurrently targets the WNT/β-catenin pathway and alleviates stemness-related features in CRPC.

The WNT/β-catenin pathway is known to be modulated by ADT in advanced-stage PCa patients and confers androgen-independent cell survival (41). The activation of the WNT/β-catenin pathway configures the enzalutamide resistance, and its inhibition resensitizes PCa cells to AR-targeted therapy (42). Additionally, the CRPC patients with increased β-catenin activity exhibited low AR levels and *vice versa* (40), and reciprocal inhibition of β-catenin and AR signaling was observed in prostate tumorigenesis (43). Similarly, our findings demonstrated that targeting KIT signaling leads to an increase in AR/REST signaling axis which in turn down-regulates SPINK1, leading to the mitigation of neuroendocrine transdifferentiation. Apart from the mutual antagonism between β-catenin and AR signaling, our study also indicates KIT signaling-mediated proteasomal degradation of AR, which might confer enzalutamide resistance (**Fig. 8)**. Overall, our findings that inhibition of KIT signaling coaxed AR/REST transactivation is of clinical relevance for CRPC patients, who display activated WNT/β-catenin pathway and therapy resistance.

**Figure 8.**
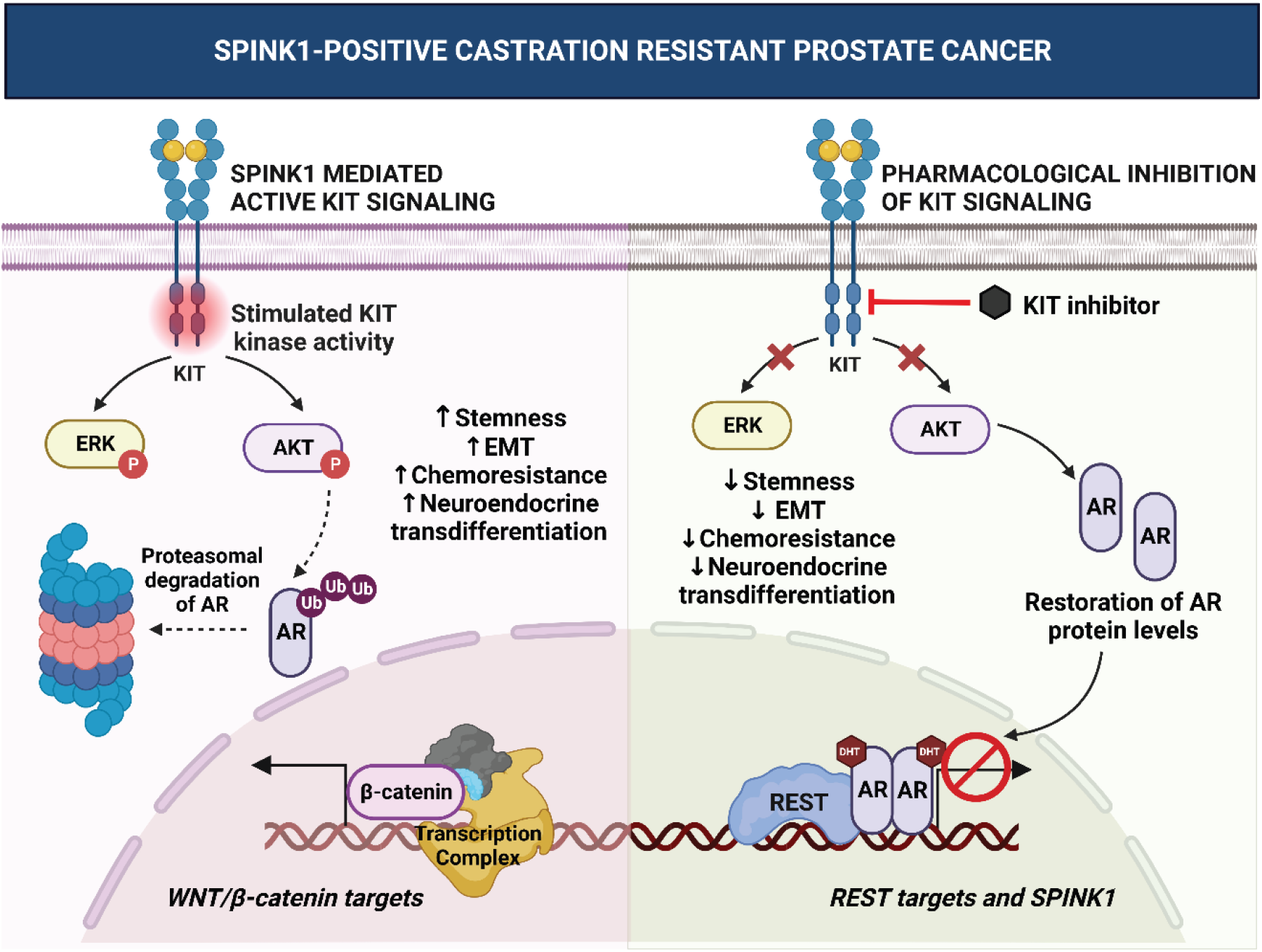
Schematic representation of KIT signaling in SPINK1-positive castration-resistant prostate cancer. SPINK1-induced KIT kinase activity triggers downstream MAPK and AKT signaling. The phosphorylated AKT might result in the proteasomal degradation of AR. The AR^low^/ KIT-positive cells transactivate β-catenin and upregulate WNT/β-catenin target genes (left panel). While pharmacological inhibition of KIT signaling restores AR levels, modulating AR/REST transcriptional activity, which downregulates SPINK1 and other REST target genes (right panel).

Imatinib, the most commonly used type III receptor tyrosine kinase inhibitor, has not delivered any favorable clinical response either as a single agent or in combination therapy for PCa patients (44-46). Most of these trials have evaluated Imatinib’s potential as a PDGFR inhibitor due to its high specificity. Therefore, we traversed for a more potent type III receptor tyrosine kinase inhibitor, namely, pexidartinib (PLX3397), which has ten to hundred times more selectivity for KIT and CSF1R than other related kinases (47). Our preclinical mice study indicates the therapeutic potential of pexidartinib in abrogating SPINK1-positive CRPC tumor progression. The systemic administration of pexidartinib with AR antagonists may yield better clinical outcomes, particularly in CRPC cases with bone metastasis. Collectively, our findings provide a compelling rationale for therapeutic targeting of KIT signaling either alone or in combination with the mainstay ADT for advanced-stage PCa patients.

## STAR Methods

### Cell lines and cell culture

The prostate cancer (LNCaP, VCaP, 22RV1, DU145, PC3, NCIH660), the benign prostatic epithelial (RWPE1 and PNT2), and the human epithelial (HEK293FT) cell lines were procured from the American Type Cell Culture (ATCC). They were cultured in their recommended media supplemented with 10% fetal bovine serum (FBS) (Thermo Fisher Scientific) and 0.5% Penicillin-Streptomycin (Pen-Strep) (Thermo Fisher Scientific) following the ATCC guidelines. The enzalutamide-resistant 42D^ENZR^ cell line was generously provided by Prof. Amina Zoubeidi and was cultured as previously described (48). The authentication of cell lines was performed using short tandem repeat (STR) profiling at the Lifecode Technologies Private Limited, Bangalore and DNA Forensics Laboratory, New Delhi. The cell lines were routinely monitored for any Mycoplasma contamination using the PlasmoTest mycoplasma detection kit (InvivoGen).

### Androgen deprivation and drug treatment

For androgen-independence, LNCaP cells were cultured in the RPMI media without phenol red (Gibco), supplemented with 5% charcoal stripped FBS for 2 weeks. For KITi treatment, cells were serum starved for 6hrs in the culture media without FBS and then treated with pexidartinib (MedChemExpress, HY-16749) for the 60hrs in complete culture media.

### Plasmids and lentiviral packaging

The lentiviral pGIPZ plasmids containing short-hairpin RNA (shRNA) such as shScrambled, and shSPINK1 were purchased from Dharmacon (Horizon Discovery Ltd.). Doxycycline-inducible KIT overexpression plasmid (pCLXEBR-pTF-cKit) (49) was a gift from Patrick Salmon (Addgene plasmid #114293). Lentiviral packaging was done using the ViraPower™ Lentiviral Expression Systems (Invitrogen) following the manufacturer’s protocol. Briefly, HEK293FT cells were plated at 90% confluency and transfected with the ViraPower packaging mix (9µg) and the lentiviral plasmids (shRNA/overexpression constructs, 3µg) using Polyethylenimine (Polysciences, 23966). After 60-72hrs transfection, the lentiviral particles were harvested and stored at −80°C. To produce stable cell line, cells were infected with the lentiviral particles and polybrene (hexadimethrine bromide; 8µg/ml) (Sigma-Aldrich, 107689). After 24hrs of infection, the culture media was changed and the shRNA stable cells were selected in puromycin (Sigma-Aldrich, P8833) and Dox inducible KIT overexpressing cells in blasticidine (Sigma-Aldrich, 15205).

### Proteome and phosphoproteome profiling

Cell lysis and protein estimation− 22RV1-shSPINK1 and 22RV1-shSCRM cells were washed with 1X phosphate buffered saline (PBS) twice and lysed in hot boiling SDS lysis buffer (5% SDS, 50mM Tris pH 8.5). The lysate was immediately heated for 5min and sonicated; then centrifuged at 18000 rpm for 15min. The supernatant was collected and the protein estimation was done using BCA. 5mg/ml concentrated protein lysate was prepared for each sample and stored in −80°C till further processing.

Sample preparation for mass spectrometry− 1mg of protein sample was used for digestion; reduced with 5mM tris(2-carboxyethyl)phosphine (TCEP) and alkylated with 50mM iodoacetamide and further digested with Trypsin (1:50) for 16hrs at 37°C. Salt was removed using C18 silica cartridge and digested peptides were dried using a speed vac and resuspended in buffer A (2% acetonitrile, 0.1% formic acid). 500µg peptides were used for phosphopeptides enrichment, while 50µg of protein sample was used for digestion in proteome profiling.

Mass spectrometric analysis− All the processed samples were subjected to an EASY-nLC 1000 Liquid Chromatograph (Thermo Fisher Scientific) system coupled with an Orbitrap Exploris™ (Thermo Fisher Scientific) Mass Spectrometer. 1µg of peptide sample were loaded on C18 column 15cm, 3.0μm Acclaim PepMap (Thermo Fisher Scientific) and separated with 0–40% gradient of buffer B (80% acetonitrile, 0.1% formic acid) at 300nl/min flow rate); LC gradients were run for 110mins. MS1 spectra was acquired with (Max IT=60ms, AGQ target=300%; RF Lens=70%; R=60K, mass range=375-1500; Profile data). Dynamic exclusion was employed for 30s and MS2 spectra was acquired for top 20 peptides with MS2 (Max IT=60ms, R=15K, AGC target 100%). For proteome profiling, MS1 spectra was acquired with (Max IT=25ms, AGQ target=300%; RF Lens=70%; R=60K, mass range=375-1500; profile data).

Data processing− All samples were processed and raw files containing mass/charge values were created for each sample. These raw files were analyzed against UniProt Human Proteome Reference through Proteome Discoverer Software (v2.5, Thermo Fisher Scientific). For SEQUEST and MS Amanda search, the precursor (10 ppm) and fragment mass tolerance (0.02 Da) were set. The enzyme specificity for trypsin/P (cleavage at the C-terminus of “K/R: unless followed by “P”) and maximum two missed cleavages were considered. Carbamidomethyl at cysteine as fixed; and phosphorylation at S, T and Y, oxidation at methionine, and acetylation at N-terminus both were set as variable modifications. The 0.01 false discovery rate (FDR) was considered for both peptide spectrum match and protein. For proteome profiling, phosphorylation at S, T and Y were not considered as variable modifications.

### Immunoprecipitation followed by mass spectrometry

Dox-induced 22RV1-KIT cells were washed with ice-cold PBS twice and lysed with ice-cold lysis buffer (50mM Tris-HCl pH-8.0, 100mM NaCl, 2% NP-40, 2% Triton X-100, 2mM EDTA). The cell lysate was kept on rotation for 60 min at 4°C and then precleared with 100µL Protein A/G PLUS-Agarose bead slurry for 30 min at 4°C. 10% of the precleared lysate was saved as input and the remaining is divided into two equal halves: one with KIT (CST) and the other with Rabbit IgG Isotype control (Invitrogen), incubate them with rotation overnight at 4°C. Protein A/G PLUS-Agarose were washed in the lysis buffer, then added to KIT and IgG containing lysates, and incubated for 4hrs with rotation at 4°C. The bead-antibody-protein complexes were washed thrice in wash buffer (10mM Tris-HCl pH-7.4, 150mM NaCl, 1% Triton X-100, 2mM EDTA). Elution was done by heating the bead-antibody-protein complex in 2X SDS loading dye without reducing agent for 10min at 75°C.

Mass spectrometry analysis− 50µg of protein was used for trypsin digestion and the sample preparation and processing for mass spectrometry was done similar to the 22RV1- shSPINK1/22RV1-shSCRM proteome profiling. MS1 spectra was acquired with (Max IT=25ms, AGQ target=300%; RF Lens=70%; R=60K, mass range=375-1500; profile data). Dynamic exclusion was employed for 30s and MS2 spectra was acquired for top 12 peptides with MS2 (Max IT=22ms, R=15K, AGC target 200%). The RAW files generated from IP-MS samples were analyzed against the UniProt Human Proteome Reference through Proteome Discoverer Software (v2.5, Thermo Fisher Scientific). For SEQUEST and MS Amanda search, the precursor (10ppm) and fragment mass tolerances (0.02 Da) were set. The enzyme specificity for trypsin/P (cleavage at the C-terminus of “K/R: unless followed by “P”) was set. Carbamidomethyl at cysteine as fixed and oxidation at methionine and acetylation at N-terminus were set as variable modifications. The 0.01 FDR was considered for both peptide spectrum match and protein.

### Cytotoxicity assay

To evaluate the half-maximal inhibitory concentration (IC50) of pexidartinib (KITi), 22RV1 (3 × 10^3^) cells were seeded in 96-well culture dishes and treated with different concentrations of KITi for 48hrs. After treatment, Cell Proliferation Reagent WST-1 (Roche) was added and the IC50 was determined following the manufacturer’s protocol.

### Cell proliferation and viability assay

22RV1/ 42D^ENZR^ (3 × 10^3^) cells were seeded in 96-well culture dishes and treated with different concentrations of pexidartinib (KITi) against DMSO (CTL) in the recommended complete media and incubated for the specified time points (up to 4 days). KITi/CTL containing media was changed after every 48hrs. At the end-point, Cell Proliferation Reagent WST-1 (Roche) was added and cell viability was determined following the manufacturer’s protocol.

### Foci formation assay

22RV1/ 42D^ENZR^ (2 × 10^3^) cells were plated in 6-well culture dishes along with the recommended media and 5% fetal bovine serum (Invitrogen). The KITi treatment was started after 2 days of cell plating and media was changed after every 48hrs. The assay was terminated after 2 weeks and the foci were fixed with 4% paraformaldehyde (in 1X PBS) and stained with crystal violet solution (0.05% w/v, 20% ethanol and 1X PBS). For quantitation, 10% glacial acetic acid was used for destaining and the crystal violet absorption was measured at 550nm. The representative images were captured using Leica DFC310 FX microscope (Leica Microsystems).

### Tumorsphere assay

22RV1 (1 × 10^4^) cells were seeded in Ultra-Low Attachment 6-well culture dishes in DMEM- F12 media (1:1, Invitrogen) along with EGF (20 ng/ml, Invitrogen), FGF (20 ng/ml, Invitrogen) and B27 (1X, Invitrogen) as previously described. After every 2 days, tumorspheres were collected, disintegrated into single cell suspension and re-plated in fresh media along with KITi/CTL. The tumorspheres were harvested after two weeks and the tumorsphere formation efficiency was determined by evaluating the number of spheres >50µm in diameter and the area of tumorspheres were calculated using ImageJ. The representative images were captured using Axio Observer Z1 inverted microscope (Carl Zeiss).

### Flow cytometry analysis

For stem cell surface marker staining, KITi/CTL treated 22RV1 (1 × 10^6^) cells were stained with 1:50 dilution of CD44-PE antibody (Miltenyi Biotec, 130-113-904) and incubated for 1hr at 4°C. The stained population of cells were analyzed by flow cytometry and live cells were gated using forward scatter (FSC) and side scatter (SSC) dot plot. The cells positive for CD44- PE staining were analyzed relative to the respective IgG isotype control. About 1 × 10^5^ events were acquired for each sample on BD FACS Melody™ Cell Sorter and analyzed with FlowJo version 10.7.

### Immunoblot analysis

Cells were washed in ice-cold PBS and protein was extracted using radioimmunoprecipitation assay (RIPA) lysis buffer, supplemented with Protease Inhibitor Cocktail (Genetix) and Phosphatase Inhibitor Cocktail Set-II (Calbiochem). The samples were prepared in Laemmli sample buffer, resolved on SDS-PAGE and transferred to polyvinylidene difluoride (PVDF) membrane (PALL). The membrane was blocked using 5% non-fat dry milk in tris-buffered saline with 0.1% Tween 20 (TBST) for 1 hr at room temperature, and incubated overnight at 4°C with the respective primary antibody: 1:1000 diluted KIT (CST, 37805), 1:1000 diluted phospho-KIT (CST, 3073), 1:3000 diluted β-catenin (CST, 8480), 1:1000 diluted CD44 (CST, 3570), 1:1000 diluted phospho-Akt (CST, 13038), 1:1000 diluted total-AKT (CST, 9272), 1:1000 diluted phospho-ERK (CST, 4377), 1:1000 diluted total-ERK (CST, 4695), 1:1000 diluted AR (CST, 5153), 1:2000 diluted REST (Abcam, ab75785), 1:1000 diluted β3-tubulin (CST, 5568), 1:500 diluted SPINK1 (R&D Systems, MAB7496-SP), and 1:5000 diluted β- actin (Abcam, ab6276). The blots were then washed thrice in 1X TBST buffer and incubated with the respective horseradish peroxidase-conjugated anti-rabbit/ anti-mouse antibody (Jackson ImmunoResearch, 711-035-150 or 711-035-152) for 2hrs at room temperature. The blots were then washed thrice in TBST, incubated with SuperSignal™ West Chemiluminescent Substrate (Thermo Fisher Scientific) and the signals were captured using X-ray films.

### Immunofluorescence staining

Cells were cultured in the specified conditions on 12mm coverslips in the 24-well culture dishes. The cells were fixed with 4% paraformaldehyde in PBS, washed with PBS, and permeabilized using PBS with 0.3% Triton X-100 for 10 min. The blocking was done using 5% normal goat serum in PBS along with 0.05% Tween 20 (PBST) for 2 hrs at room temperature. The cells were then incubated with 1:100 diluted CD44 (CST, 3570) primary antibody in PBST overnight at 4°C. Cells were washed thrice using PBST, followed by incubation with 1:600 diluted Alexa Fluor 555 conjugated anti-mouse secondary antibody (CST, 4409) in PBST for 1hr at room temperature. Cells were washed in PBST and then stained with DAPI (Sigma-Aldrich). The coverslips were mounted with VECTASHIELD antifade mounting medium (Vector laboratories) on glass slides and sealed with nail polish to prevent drying. The representative images were captured using Axio Observer Z1 motorized inverted fluorescence microscope (Carl Zeiss).

### Quantitative-PCR (QPCR) analysis

The total RNA was extracted using RNAiso Plus (Takara) and 1µg of total RNA was used as template for cDNA synthesis using First Strand cDNA Synthesis Kit (Genetix) following the manufacturer’s protocol. For qPCR analysis, each reaction was set up using cDNA template, SYBR Green qPCR Master Mix (Genetix) and the respective primer set (Table S2). The qPCR reactions were performed in triplicates on StepOne Real-Time PCR System (Applied Biosystems) and the relative target gene expression was determined using the ΔΔCt method.

### Prostate cancer patient specimens

The tissue microarrays (TMAs) comprising of prostate cancer patients were obtained from the Department of Pathology, Henry Ford Health System, Detroit, MI. The TMAs contain PCa cases with radical prostatectomy, and most of them were localized cancer cases and some with lymph node metastases. All specimens were collected after receiving patient’s informed consent and Institutional Review Board approval following the ethical principles instated by Declaration of Helsinki. The TMAs were stained for KIT and SPINK1 using IHC.

### Immunohistochemistry

Slides were incubated at 60°C for at least 2hrs and then placed in EnVision FLEX Target Retrieval Solution, either low pH (Agilent DAKO, K800521-2) or high pH (Agilent DAKO, K800421-2) in a PT Link instrument (Agilent DAKO, PT200) at 75⁰C, heated to 97⁰C for 20min, and then cooled to 75⁰C. Next, slides were washed in 1X EnVision FLEX Wash Buffer (Agilent DAKO, K800721-2) for 5min. Slides were treated with Peroxidazed 1 (Biocare Medical, PX968M) for 5min and Background Punisher (Biocare Medical, BP974L) for 10min with a wash of 1X EnVision FLEX Wash Buffer for 5min after each step. 1:100 diluted SPINK1 [4D4] (Novus Biologicals, H00006690-M01), 1:20 diluted CD117/KIT (DAKO, A4502), 1:25 diluted CD44 (DAKO, M7082), Ki67 [MIB-1] (DAKO, IR626) in EnVision FLEX Antibody Diluent (Agilent DAKO, K800621-2) was added to each slide and incubated overnight at 4⁰C. Slides were then washed in 1X EnVision Wash Buffer for 5min and incubated in either Mach2 Doublestain 1 (Biocare Medical, MRCT523L) (rabbit) or Mach2 Doublestain 2 (Biocare Medical, MRCT525L) (mouse) for 30min at room temperature in a humidifying chamber. Next, slides were rinsed in 1X EnVision Wash Buffer thrice for 5min each and treated with a Betazoid DAB solution (Biocare Medical, BDB2004L) for 5 mins. Slides were rinsed twice in distilled water, and treated with EnVision FLEX Hematoxylin (Agilent DAKO, K800821-2) for 5min. After several rinses times in tap water and drying, slides were dipped in xylene approximately 15 times. EcoMount (Biocare Medical, EM897L) was added to each slide, which was then cover slipped.

IHC staining analysis− The scoring for SPINK1 IHC staining was considered as either positive or negative as has been mentioned previously (50). The scoring for KIT IHC staining was categorised as either: high, medium, low, or negative, based upon the intensity. For the xenograft model, 5 fields were randomly selected from each tumor tissue and the IHC scoring was categorised as follows: 0 (negative), 1 (weak), 2 (moderate), and 3 (strong), based upon the intensity.

### Mice xenograft studies

The immunodeficient NOD/SCID mice (NOD.Cg-Prkdc^Scid^/J) were procured from The Jackson Laboratory and were maintained as per their recommended guidelines. For xenograft experiments, 5-6 weeks old NOD/SCID male mice were intraperitoneally injected with anesthetic cocktail of ketamine (50 mg/kg) and xylazine (5 mg/kg). 22RV1 (3 × 10^6^) cells with 30% Matrigel (Corning) resuspended in 100µl saline were subcutaneously implanted in the dorsal flanks of mice. Mice were routinely monitored for tumor growth and once the tumor volume reached ∼75 mm^3^, mice were randomized into 2 groups (n = 5 each). The treatment groups were orally administered with either pexidartinib (KITi) (MedChemExpress, HY- 16749; 50 mg/kg mice body weight) or control [5% dimethyl sulfoxide (DMSO)] along with 30% polyethylene glycol-400 (PEG-400) and 65% saline, five times a week. The tumor growth was measured every alternate day using Vernier caliper, and tumor volumes were measured using the formula (π/6) (W^2^×L), (W=width and L=length). After three weeks of drug treatment, the mice were euthanized and tumors, lungs, and bone marrows were harvested. Tumors were fixed in 10% neutral buffered formalin (NBF), embedded in paraffin and subjected to IHC staining for the specific markers. For spontaneous metastasis, genomic DNA (gDNA) from lungs and bone marrow was extracted and subjected to TaqMan assay using TaqMan TAMRA probe (Applied Biosystems, 450025), TaqMan Universal PCR Master Mix (Applied Biosystems, 4304437), and Human *Alu* primers (Table S2). The relative quantitation for the number of cells metastasized was evaluated based on a standard curve for serially diluted human gDNA spiked with mouse gDNA as previously described (30).

For experimental bone metastasis, 22RV1 (4 × 10^5^) cells resuspended in 20µl saline were implanted via intratibial injection with the 26-gauge needle in anaesthetized 5-6 weeks old NOD/SCID male mice. Post-implantation, mice were administered with Piroxicam (3mg/kg) via intramuscular injection to relieve pain. Subsequently, mice were randomized into 2 groups (n = 5 each) and were administered with either pexidartinib (KITi) (50 mg/kg) or control (5% DMSO) along with 30% PEG-400 and 65% saline via oral gavage, five times a week. After 3 weeks of drug treatment, mice were X-ray scanned using InAlyzer (MEDIKORS Inc.), euthanized and the tibiae were harvested for micro-CT analysis.

All procedures in the mice xenograft studies were implemented in accordance with the guidelines of Institutional Animal Ethics Committee of the Indian Institute of Technology Kanpur, India and approved by the Committee for the Purpose of Control and Supervision of Experiments on Animals (CPCSEA), Ministry of Environment, Forest and Climate Change, Govt. of India.

### Micro-computed tomography (µCT)

The scanning analysis of the tibiae were performed using the micro-CT system SkyScan (Bruker, 1172). The parameters considered for scanning were as follows: 7 µm resolution, 48 kV voltage, 204µA current and 0.5µm Al filter. The pixel settings were medium with 48 kV voltage and 204µA current along with 0.4 rotation step; each sample was scanned in 28min. The CTVox software was used for 3D visualization and CTAn software for 3D image processing and bone morphometric analysis.

### In silico analysis

The cut-off for differential expression of proteins and phosphopeptides were considered to be: log_2_Fold change <−0.6 for downregulated and >0.6 for upregulated, P-value <0.05. The kinase- substrate enrichment analysis was performed using KSEA App (19). The enrichment of biological processes was done using DAVID functional annotation tool (21). The correlation of *KIT*, *SPINK1* and *INSR* expression with AR signaling score and NEPC score were analyzed for Metastatic Prostate Adenocarcinoma (SU2C/PCF Dream Team, PNAS 2019 (51)) using cBioPortal. The heatmaps were generated using pheatmap (version 1.0.12) package, volcano plots using ggplot2 package and pathway enrichment using pathfindR (20) in R 4.2.1.

## Statistical Analysis

Statistical significance was measured using GraphPad Prism with the following tests: One-way ANOVA, two-way ANOVA, unpaired two-tailed Student’s t-test along with the multiple comparison analysis or otherwise indicated in the respective figure legend. The comparison between the groups was considered significant if P-value <0.05; and indicated as follows: *P < 0.05 and **P < 0.001. All the experiments were conducted in replicates and the error bars denote the standard error of mean (SEM) of at least three independent replicates.

## Acknowledgements

The illustrations in the Supplementary Figure S1 and Figure 8 are created using BioRender (https://biorender.com/). We are thankful to Prof. Jonaki Sen and Prof. Pradip Sinha for extending the use of microscope facility, Prof. Ashok Kumar for micro-CT SkyScan system and Prof. Amitabha Bandyopadhyay for CTAn software. We also acknowledge Shraddha Singh and Irfan Qayoom for their assistance with micro-CT data acquisition and visualization.

## Author Contributions

N.M. and B.A. conceptualized the study. N.M. and B.A. designed the methodology, N.M. conducted the in vitro experiments and N.M. and B.A. performed the mice xenograft studies. N.M. visualized the proteome and phosphoproteome data, gene expression datasets, and performed data analysis. U.K.K. performed Immunoprecipitation, U.K.K. and A.G. performed TaqMan-based assay and micro-CT data analysis. N.P. and N.G. assembled and provided the PCa tissue microarrays and related clinical information. S.C. performed the immunohistochemical staining. N.M. and B.A. wrote the research manuscript. B.A. supervised the overall project.

## Data Availability

The proteome and phosphoproteome profiles of the 22RV1-shSCRM and 22RV1-shSPINK1 cells were deposited in the PRIDE consortium. The publicly available gene expression datasets were downloaded from cBioPortal, namely: Metastatic Prostate Adenocarcinoma (SU2C/PCF Dream Team, PNAS 2019) and Neuroendocrine Prostate Cancer (Multi-Institute, Nat Med 2016). Other datasets used in the study were retrieved from NCBI GEO, such as GSE48403 for pre- and post-ADT treated PCa specimens, GSE112786 for NEPC organoids and GSE183199 for Enzalutamide-resistant cell line models.

