## Supplementary Figure 1 to 8 for "An integrative proteomics approach identifies tyrosine kinase KIT as a novel therapeutic target for SPINK1-positive prostate cancer"

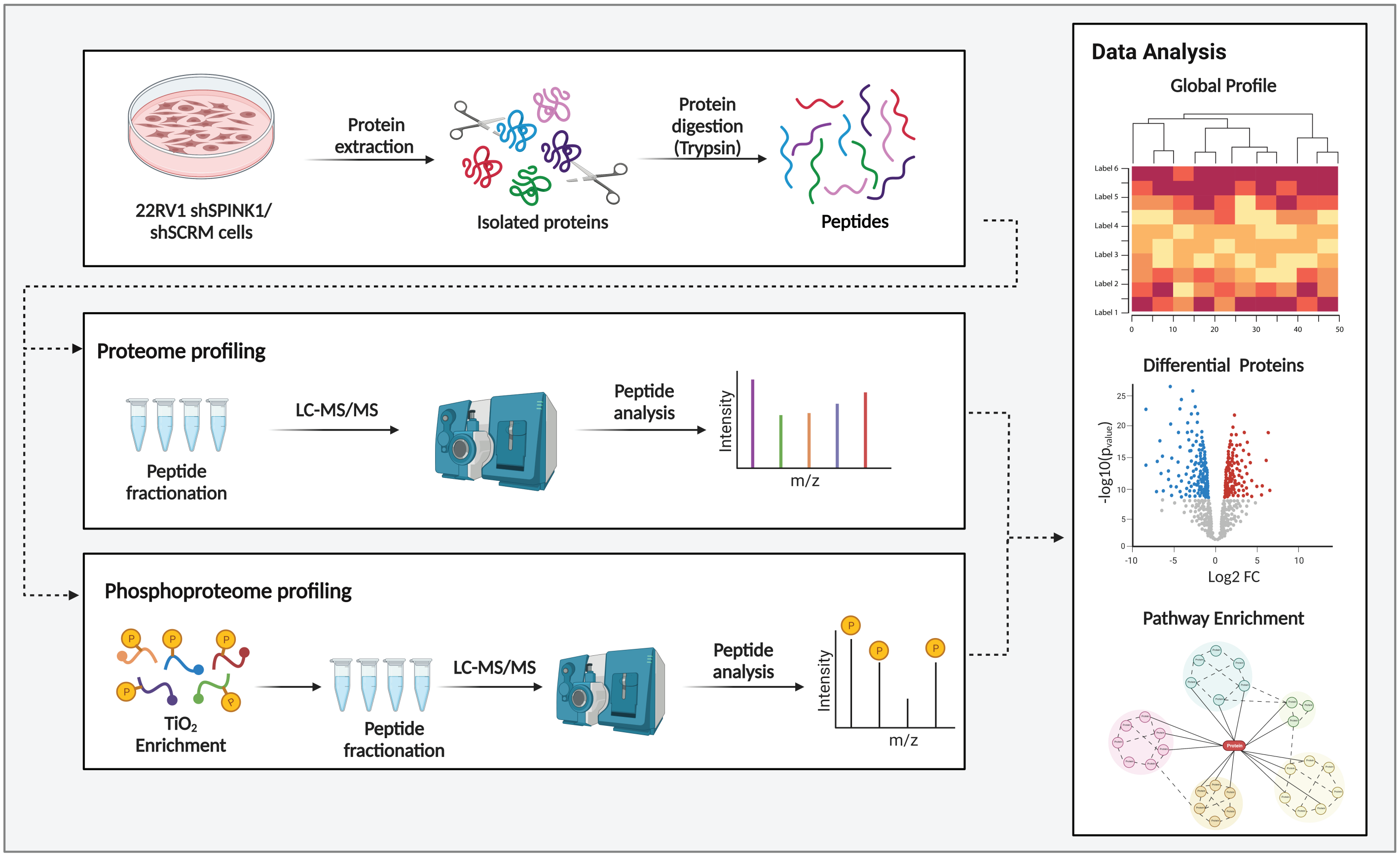


**Figure S1. Quantitative proteome and phosphoproteome profiling of the SPINK1-positive prostate cancer cells.** The proteins extracted from the 22RV1-shSCRM and 22RV1-shSPINK1 cells after lysis were digested using Trypsin to give small peptides. These peptides were fractionated and analyzed using Liquid Chromatography coupled Tandem Mass Spectrometry (LC-MS/MS). The phosphoproteome analysis was done in the similar way except for phosphopeptide enrichment using titanium dioxide ($TiO_2$) beads. Data analysis was done to enrich differential proteins and phosphoproteins and pathways were mapped.

### Supplementary Figure 2

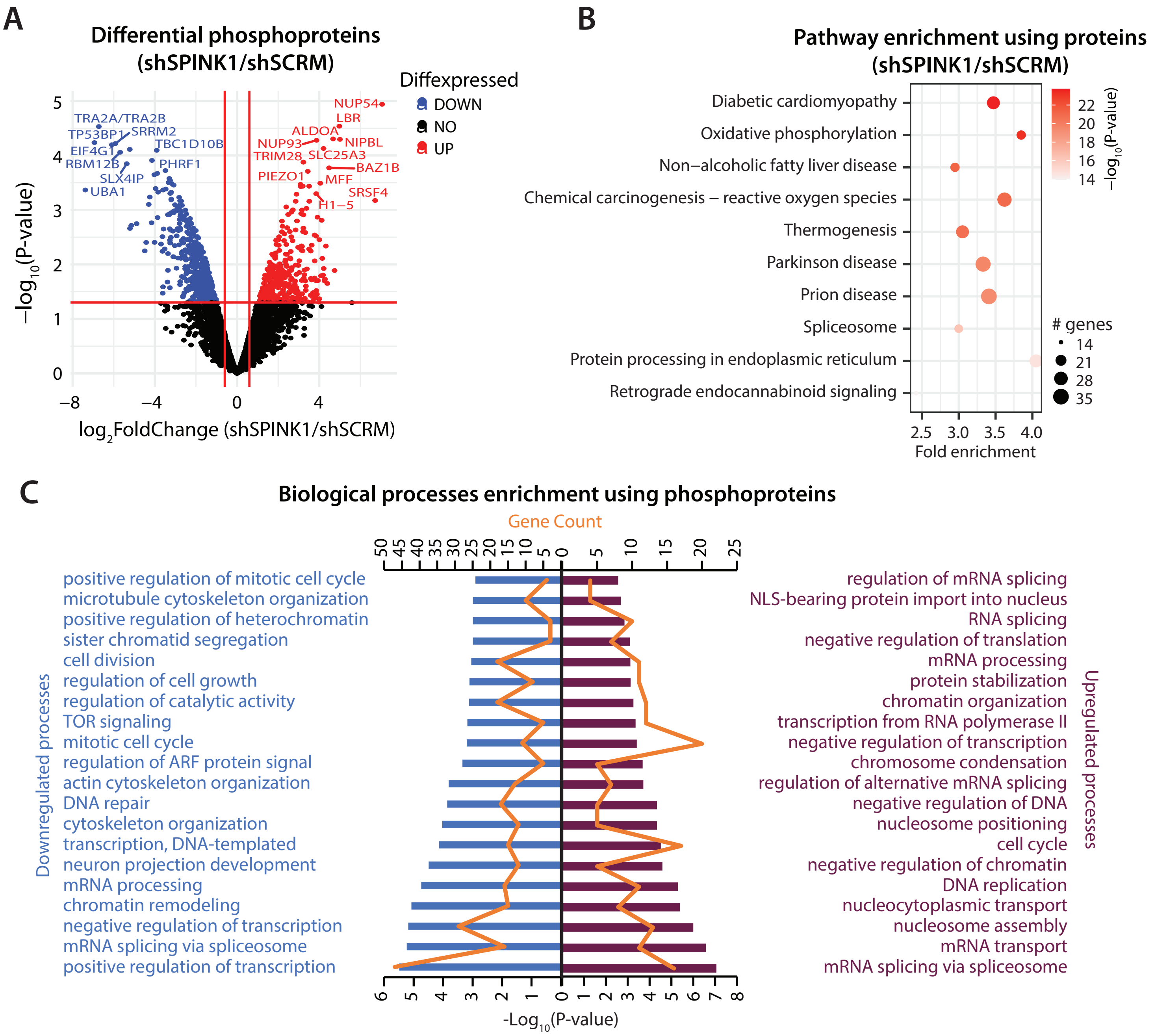


**Figure S2. Pathway enrichment analysis for SPINK1-silenced 22RV1 prostate cancer cells.** (**A**) Volcano plot showing the differentially enriched phosphoproteins in 22RV1-shSPINK1 versus 22RV1-shSCRM cells; blue dots are downregulated and red dots are upregulated phosphoproteins in 22RV1-shSPINK1 versus 22RV1-shSCRM, whereas black dots signify no change. (**B**) Pathway enrichment analysis for differential proteins in 22RV1-shSPINK1 versus 22RV1-shSCRM cells using pathfindR; the size of dot denotes the number of genes and the color symbolize the -$\log_{10}$(P-value) of the enriched pathway. (**C**) DAVID functional annotation analysis showing enriched biological processes using differential phosphoproteins; blue represents downregulated processes while magenta represents upregulated processes in 22RV1-shSPINK1 versus 22RV1-shSCRM cells. The bar denotes -log10(P-value) and the line represents gene count.

### Supplementary Figure 3

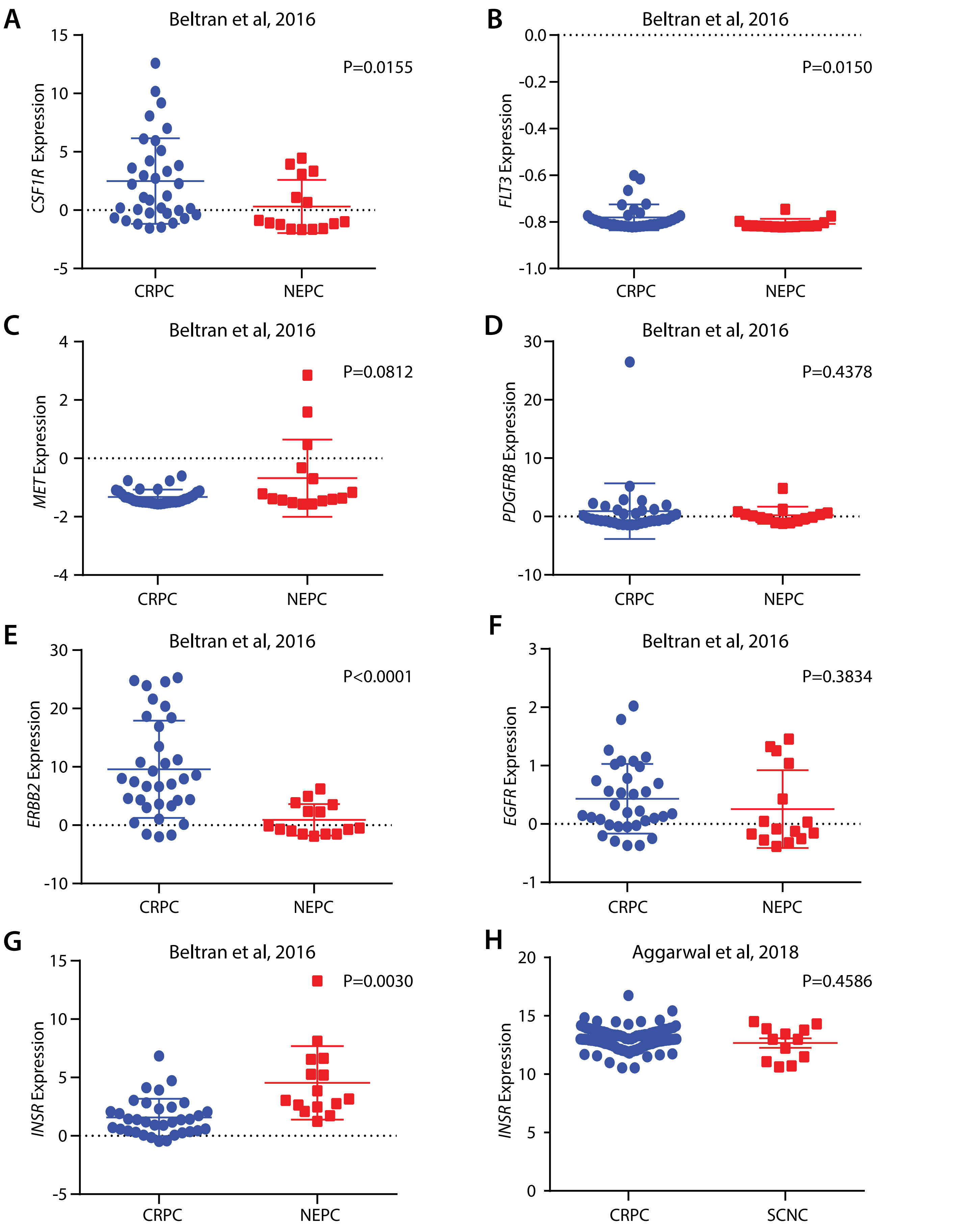


**Figure S3. Expression of receptor tyrosine kinases in advanced-stage PCa.** The mRNA expression in Beltran et al, 2016 dataset for (**A**) *CSF1R*, (**B**) *FLT3*, (**C**) *MET*, (**D**) *PDGFRB*, (**E**) *ERBB2*, (**F**) *EGFR* and (**G**) *INSR* receptor tyrosine kinases. (**H**) The mRNA expression of *INSR* in Aggarwal et al, 2018 dataset. P value was calculated using Unpaired Students' t-test with Welch's correction.

### Supplementary Figure 4

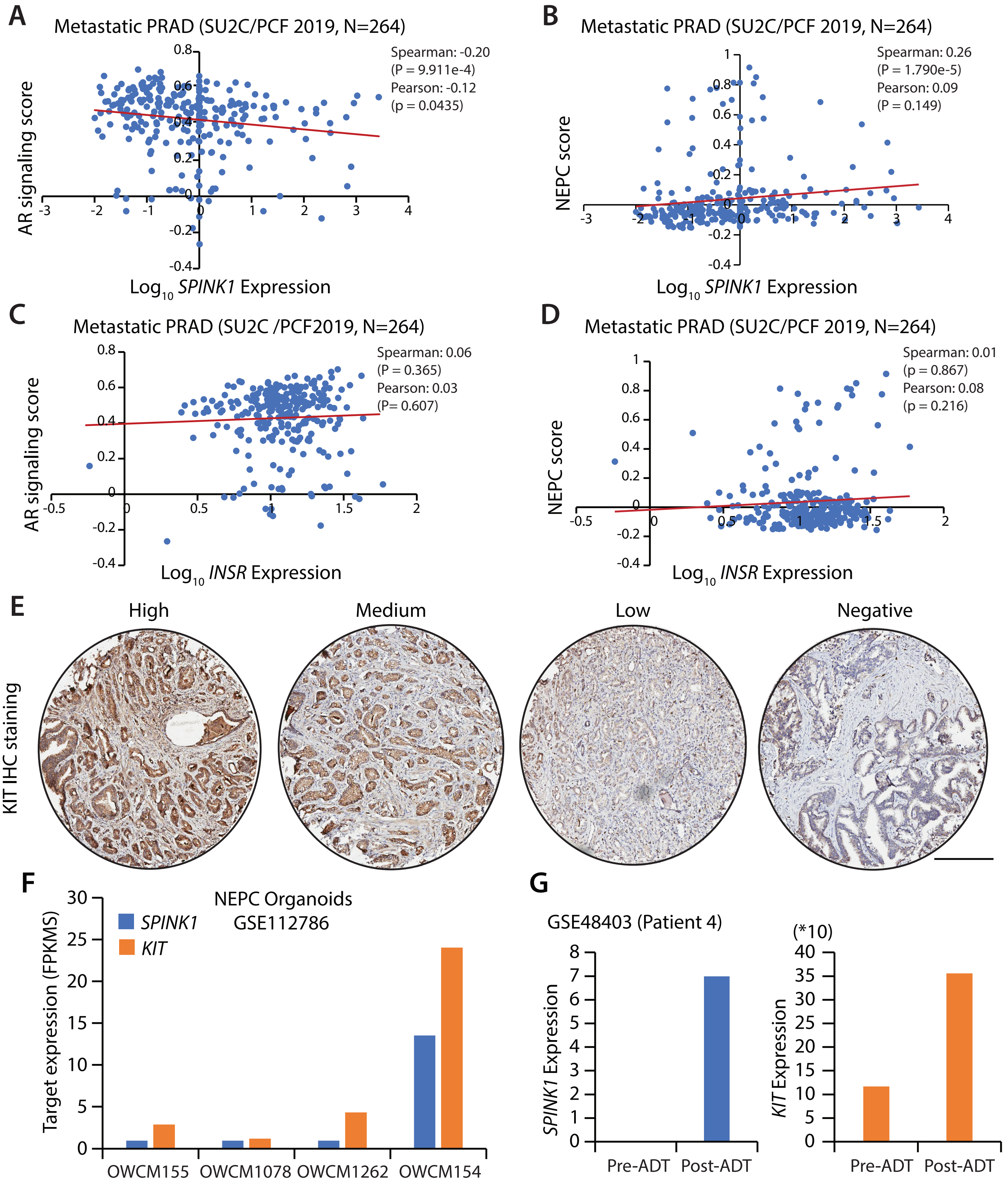


**Figure S4. KIT is inversely correlated with AR signaling similar to SPINK1.** (**A**) Scatter plot showing correlation of *SPINK1* mRNA expression (FPKM polyA) ($log_{10}$(value + 1)) with AR signaling score in metastatic prostate adenocarcinoma (SU2C/PCF 2019) dataset. Coefficient value for both Spearman and Pearson's correlation along with P value are depicted in the plot. (**B**) Same as in (A) except for correlation of *SPINK1* mRNA expression with NEPC score. (**C**) Same as (A) except for correlation of *INSR* mRNA expression and AR signaling score. (**D**) Same as (B) except for correlation of *INSR* mRNA expression and NEPC score. (**E**) Micrographs representing immunohistochemical (IHC) staining for KIT levels in tissue microarrays of prostate cancer specimens. IHC scoring is classified as high, medium, low and negative based on the intensity of staining and number of positive cells. Scale bar represents 300µm. (F) Bar plot depicting mRNA expression (FPKMS) of *SPINK1* and *KIT* in NEPC organoids. (G) Bar plots showing *SPINK1* and *KIT* expression (normalized read counts) in 22 weeks ADT treated metastatic PCa patient with Gleason score-8 and TNM stage-3a.

### Supplementary Figure 5

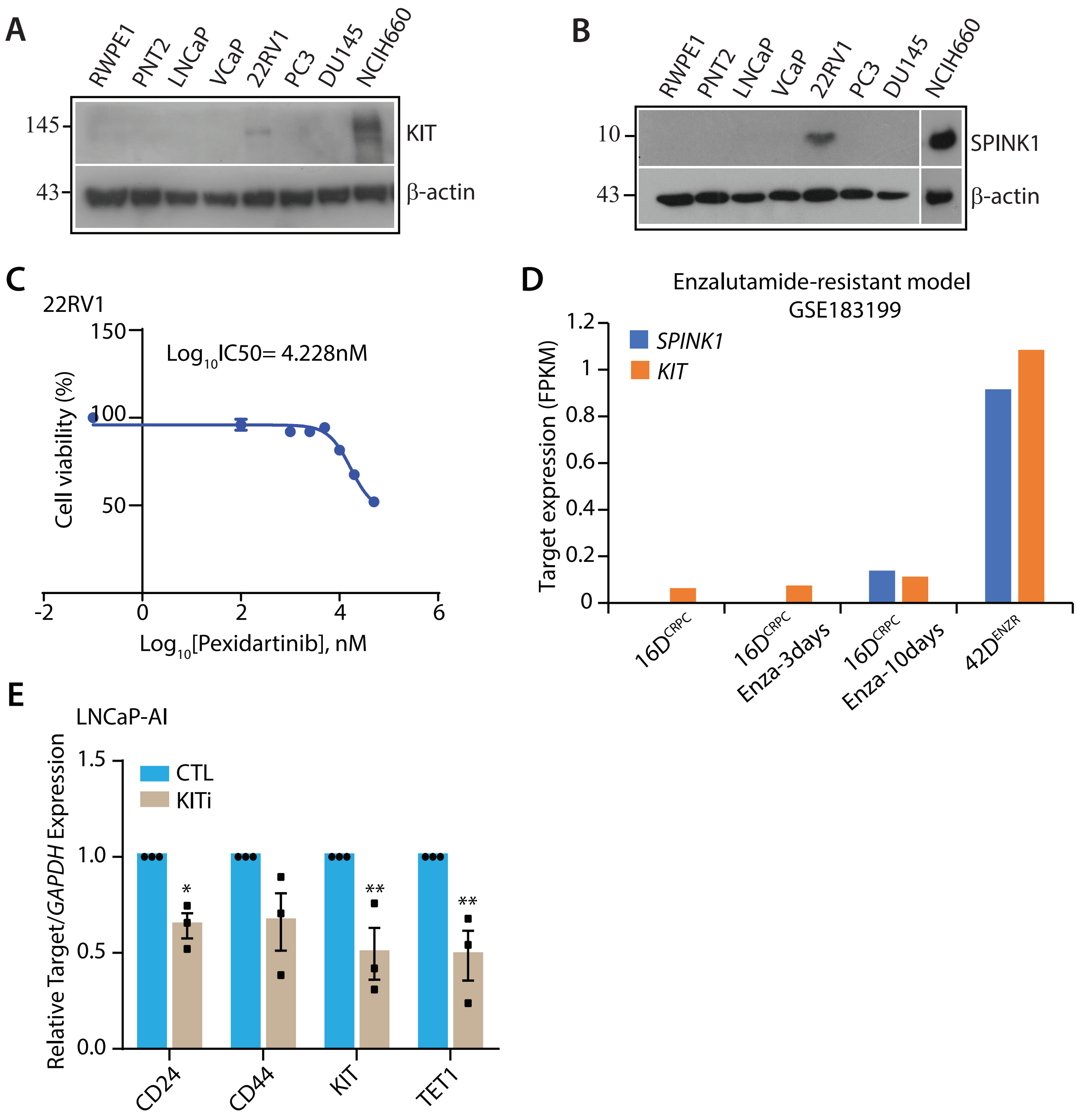


**Figure S5. KIT and SPINK1 are highly expressed in androgen-independent prostate cancer cells.** (**A**) Immunoblot showing KIT levels in a panel of benign prostate cells and prostate cancer cells. β-actin was used as loading control. (**B**) Same as (A) except for SPINK1 expression in the panel. (**C**) Line plot depicting the percent cell viability of 22RV1 cells to assess the half-maximal inhibitory concentration (IC50) of KIT tyrosine kinase inhibitor (KITi), where $\log_{10}$IC50=4.228nM (~16.9uM). (**D**) Bar plot showing mRNA expression (FPKM) of *SPINK1* and *KIT* in castration-resistant cell line model 16D$^{CRPC}$, along with enzalutaimde (Enza) treatment for 3 and 10 days, and enzalutamide-resistant cell line model 42D$^{ENZR}$. (**E**) Bar plot depicting QPCR data for the relative expression of stemness-related markers in KITi (10µM)/CTL-treated androgen-deprived LNCaP (LNCaP-AI) cells. The experiment was performed in biological triplicates (N=3); bar represents mean ± SEM and each dot represents individual value. Statistical significance was calculated using Two-way ANOVA followed by Sidak's multiple comparison test. P-value: *<0.05 and **<0.005.

### Supplementary Figure 6

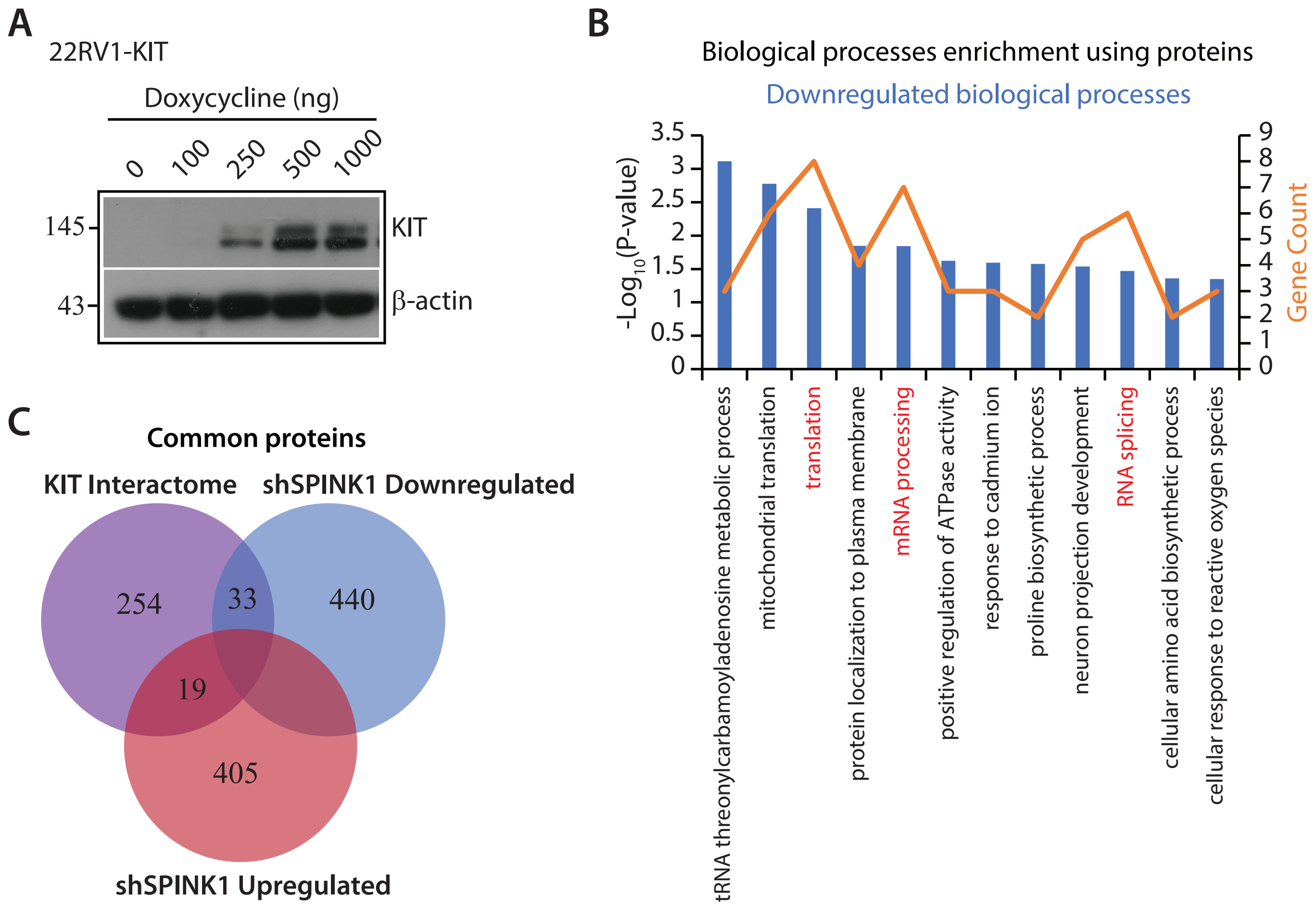


**Figure S6. KIT interactome shares SPINK1 regulated processes in prostate cancer.** (**A**) Immunoblot showing KIT levels along with different amount of doxycycline (Dox) in 22RV1-KIT cells. β-actin was used as loading control. (**B**) DAVID analysis showing enriched biological processes using downregulated proteins in 22RV1-shSPINK1 versus 22RV1-shSCRM cells. The bar denotes -$\log_{10}$(P-value) and the line represents gene count. (**C**) Venn diagram showing shared proteins among KIT interactome, downregulated in 22RV1-shSPINK1 and upregulated in 22RV1-shSPINK1 in compraison to 22RV1-shSCRM cells.

#### Supplementary Figure 7

**A** Mice tibiae with 22RV1 implantation (X-ray scan)

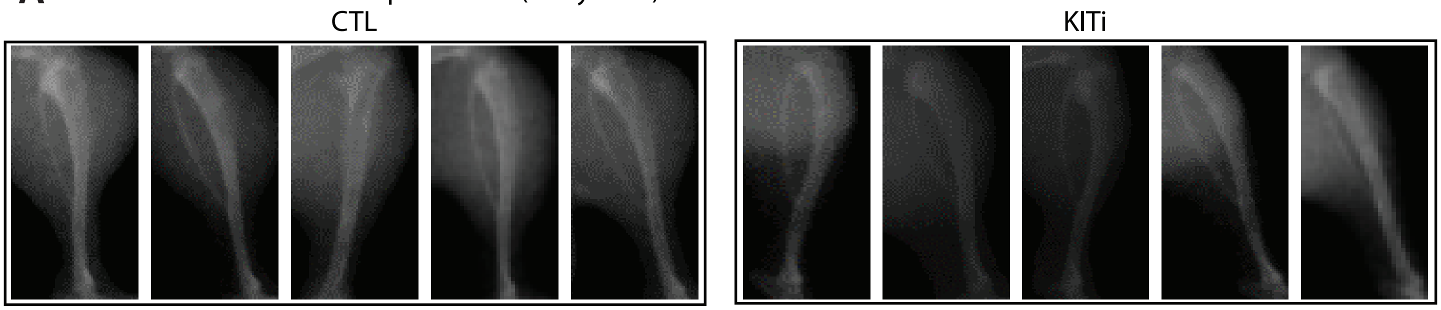


**B** Mice tibiae with 22RV1 implantation (Body compostion)

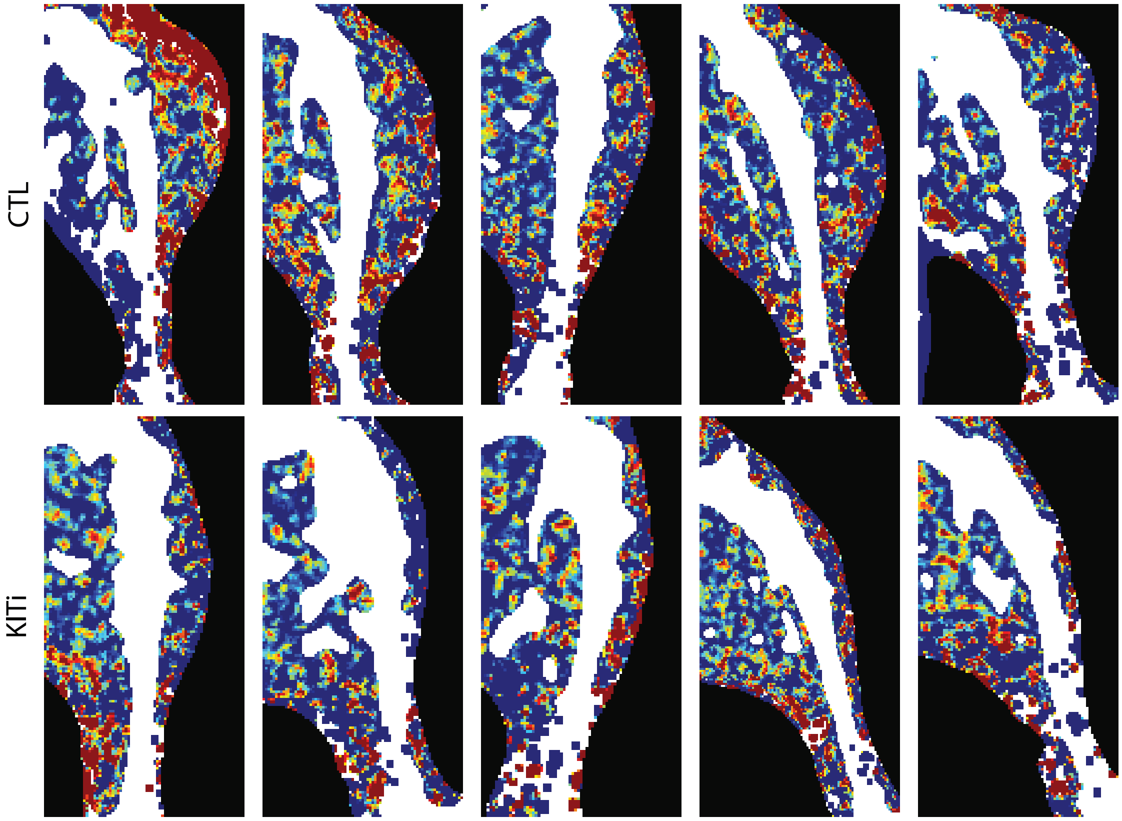


**Figure S7. KIT tyrosine kinase inhibition limits metastatic bone lesions.** (**A**) X-ray scans of the tibiae from mice implanted with 22RV1 cells via intratibial injection and administered with KITi (50mg/kg)/CTL for three weeks via oral gavage; (each group, N=5). (B) Body composition view of the tibiae same as in (A) captured via Dual Energy X-ray Absorptiometry (DEXA) system.

### Supplementary Figure 8

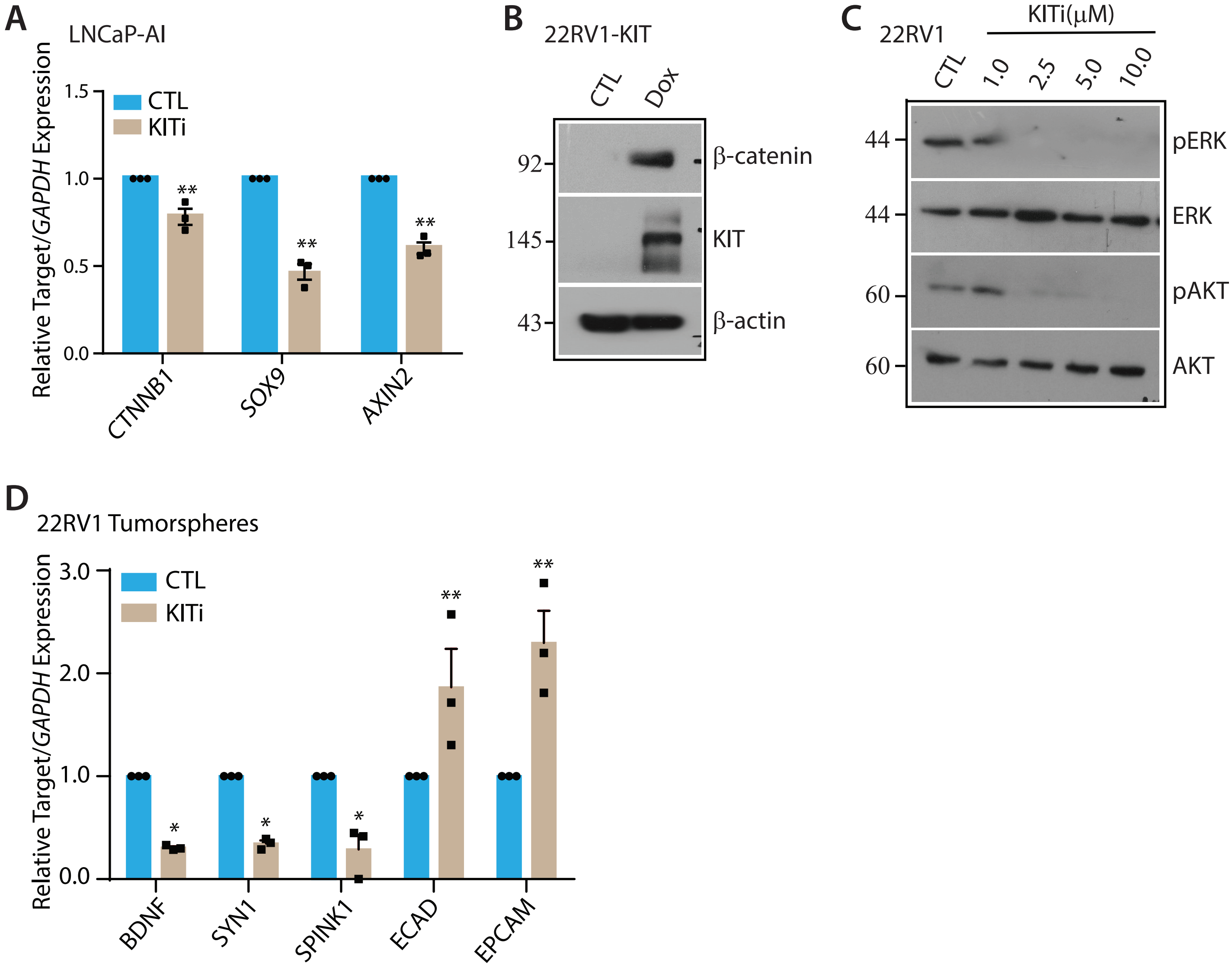


**Figure S8. KIT signaling modulates WNT/β-catenin patwhay in prostate cancer.** (**A**) Bar plot depicting QPCR data for the relative expression of WNT/β-catenin pathway genes in KITi (10μM)/CTL-treated LNCaP-AI cells. (**B**) Immunoblot showing β-catenin and KIT levels along with doxycycline (Dox) induction in 22RV1-KIT cells. β-actin was used as loading control. (**C**) Immunoblot showing expression level of phosphorylated-ERK (pERK), ERK, phosphorylated-AKT (pAKT) and AKT in 22RV1 cells treated with indicated concentrations of KITi and CTL. β-actin was used as loading control. (**D**) Bar plot showing QPCR data for the relative expression of REST target genes, SPINK1 and EMT markers in KITi (10μM)/CTL-treated 22RV1 tumorspheres. Each experiment was performed in biological triplicates (N=3); bar represents mean ± SEM and each dot represents individual value. Statistical significance was calculated using Two-way ANOVA followed by Sidak's multiple comparison test. P-value: *<0.05 and **<0.005.
